# Enlarged cortical cells and reduced cortical cell file number improve growth under suboptimal nitrogen, phosphorus and potassium availability

**DOI:** 10.1101/2020.07.06.189514

**Authors:** Xiyu Yang, Miranda Niemiec, Jonathan P. Lynch

## Abstract

Reduced cortical cell files (CCFN) and enlarged cortical cells (CCS) reduce root maintenance costs. We used *OpenSimRoot*, a functional-structural model, to test the hypothesis that larger CCS, reduced CCFN, and their interactions with root cortical aerenchyma (RCA), are useful adaptations to suboptimal soil N, P, and K availability. Interactions of CCS and CCFN with lateral root branching density (LRBD) and increased carbon availability were evaluated under limited N, P and K availability. The combination of larger CCS and reduced CCFN increases the growth of maize up to 105%, 106%, and 144%, respectively, under limited N, P, or K availability. Interactions among larger CCS, reduced CCFN, and greater RCA results in combined growth benefits of up to 135%, 132%, and 161% under limited N, P, and K levels, respectively. Under low phosphorus and potassium availability, increased LRBD approximately doubles the utility of larger CCS and reduced CCFN. The utility of larger CCS and reduced CCFN is reduced by greater C availability as may occur in future climate scenarios. Our results support the hypothesis that larger CCS, reduced CCFN, and their interactions with RCA could increase nutrient acquisition by reducing root respiration and root nutrient demand. Phene synergisms may exist between CCS, CCFN, and LRBD. Natural genetic variation in CCS and CCFN merit consideration for breeding cereal crops with improved nutrient acquisition, which is critical for global food security.

**One sentence summary:** Functional-structural modeling indicates that enlarged root cortical cells and reduced cortical cell file number decrease root maintenance cost, permitting greater soil exploration, resource capture, and plant growth under suboptimal nitrogen, phosphorus and potassium availability.

## Introduction

The development of crops with reduced fertilizer requirements is needed in global agriculture to reduce the environmental, economic, and energy costs of crop production in high-input agroecosystems and increase crop production in low-input agroecosystems (Koevoets et al., 2016; Lynch, 2019). One avenue towards this goal is via selection for root phenotypes that reduce the metabolic cost of soil exploration (Lynch, 2015). The metabolic costs of root tissues can be estimated as the investment of limiting resources, mainly carbohydrates and limiting mineral nutrients, in root growth and maintenance, and are important drivers of tolerance to edaphic stress (Chimungu et al., 2015a; Lynch, 2015; Postma and Lynch, 2011a; Saengwilai et al., 2014; Zhu et al., 2010a). The carbon cost of soil exploration includes carbon expenditure in root tissue construction and maintenance and ion uptake and assimilation (Nielsen et al., 1994, 2001). Of these, maintenance respiration is the largest carbon cost over time. Root metabolic costs at low nutrient availability are significantly greater than rates at high nutrient availability (Lambers et al., 2008; Nielsen et al., 2001). Root respiration is also a major cause of growth reduction under nutrient stress (Postma and Lynch, 2011), the consumption of carbon by root respiration can exceed 50% of daily photosynthesis under suboptimal nutrient levels (Ho et al., 2005; Lambers and Oliveira, 2020; Nielsen et al., 2001). Therefore, phenes, i.e., the basic unit of the phenotype, and phene states, i.e., the status of specific phenes (Lynch, 2011; Pieruschka and Poorter, 2012; York et al., 2013), that reduce maintenance respiration allow more internal resources to be allocated to better root establishment, thus improving crop growth under limited nutrient availability, and therefore present opportunities for the development of crops with reduced nutrient requirements (Lynch, 2015).

The “Steep, Cheap and Deep” (SCD) ideotype proposes maize root phenotypes to optimize water and N capture under limited availability of those resources (Lynch, 2013). This ideotype consists of root anatomical, architectural and physiological phenes that increase root depth, and improve the acquisition of resources from deep soil domains. By determining the proportion of respiring to non-respiring root tissue affecting the carbon and nutrient cost of tissue construction and maintenance, root anatomy regulates the metabolic cost of soil exploration and therefore is an important factor in the effects of edaphic stress on root and whole plant development (Fan et al., 2003; Jaramillo et al., 2013; Mano et al., 2006). The “topsoil foraging” ideotype for P capture (Ho et al., 2004; Lynch, 2011; Lynch and Brown, 2001; Richardson et al., 2011; Wang et al., 2010, Lynch, 2019) has been useful as a breeding goal in developing soybean and common bean cultivars that can enhance P acquisition in low phosphorus and drought environments (Burridge et al., 2019), with similar application for enhanced P acquisition in maize (Zhu et al., 2005), given that P is immobile in the soil strata, and is concentrated in the topsoil. Phene states that create a greater root surface area in the topsoil, such as shallow root angle (Lynch and Brown, 2001; Rubio et al., 2003; Zhu et al., 2005; Rangarajan et al., 2018) many hypocotyl-borne roots (Miller et al., 2003; Walk et al., 2006; Rangarajan et al., 2018), dense lateral branching (Zhu and Lynch, 2004; Jia et al., 2018), greater production of axial roots (Miguel et al., 2015, Walk et al., 2006; Rangarajan et al., 2018), RCA formation (Postma and Lynch, 2011a,b) and root hair formation (Zhu et al., 2010b; Miguel et al., 2015), have greater capacity of intercepting P, thus enhancing P uptake in the topsoil. This strategy may also be relevant to improving K acquisition in the topsoil under low K availability, as K is also relatively immobile (Lynch, 2019). Anatomical phene states that contribute to reduced metabolic cost were also found to exhibit synergism with architectural phenes (Postma and Lynch, 2011a).

The formation of root cortical aerenchyma (RCA), the enlarged intercellular spaces that form through either programmed cell death or cell separation (Evans, 2003), is generally increased in response to hypoxia (Jackson and Armstrong, 1999) and various edaphic stresses, including suboptimal availability of phosphorus, nitrogen, sulfur, and water (Bouranis et al., 2003; Drew et al., 1989; Fan et al., 2003; Konings and Verschuren, 1980; Zhu et al., 2010a; Saengwilai et al., 2014; Chimungu et al., 2015; Galindo-Castañeda et al., 2019). RCA formation alleviates the limitation of hypoxia for root respiration with improved oxygen transport (Jackson and Armstrong, 1999). The utility of RCA formation to maintain greater growth rates under various soil nutrient and drought stresses by remobilizing nutrients from the root cortex and reducing maintenance respiration has been demonstrated in several previous studies (Chimungu et al., 2015a; Fan et al., 2003; Galindo-Castañeda et al., 2019; Jaramillo et al., 2013; Postma and Lynch, 2011a; Saengwilai et al., 2014; Zhu et al., 2010a). However, the dynamic interaction between RCA and other anatomical phenes and their effects on growth requires further examination, as RCA formation reduces the proportion of root volume occupied by living cortical tissue, which is more metabolically demanding than stelar tissue (Lynch, 2013). Chimungu et al. (2014a, b) reported that reduction in the number of concentric layers of parenchyma cells in the cortex of the maize root, or cortical cell file number (CCFN), and increased volume of individual cortical parenchyma cells, or cortical cell size (CCS), could decrease the metabolic costs of root growth and maintenance, in terms of both the carbon cost of root respiration and the nutrient content of cortical tissue. In contrasting maize lines exposed to water deficit stress in controlled environments and the field, larger CCS and reduced CCFN were associated with reduced root respiration, deeper rooting, greater water capture, improved plant water status, and hence greater growth and yield (Chimungu et al., 2014a, b). However, the physiological utilities of larger cortical cells, reduced CCFN, and their interaction with RCA and root architectural phenes under nutrient deficiencies, are not known.

The utility of a root phene state under stress may be dependent on its interactions with other architectural and anatomical phenes. Phene synergism refers to the phenomenon where the combined effect of two or more phenes is greater than the additive sum of their individual effects. For example, in low P soils, common bean genotypes with long root hairs (RHL) and shallow basal root growth angle (BRGA) had three-fold greater biomass accumulation than genotypes with short root hairs and steep root angle, while only 89% greater biomass was contributed by RHL alone, and 58% by shallow BRGA alone (Miguel et al., 2015). In another study, RCA formation in lateral roots in genotypes with increased lateral root branching density had greater benefits for phosphorus acquisition (Postma and Lynch, 2011b) than the effect of RCA alone. Integration of anatomical phenes and architectural phenes of maize root systems are important for plant growth and nitrogen acquisition (York et al., 2013; York and Lynch, 2015). These potential synergisms may be useful for breeding crops with greater edaphic stress tolerance. However, interactions among phenes may also be antagonistic, i.e., the functional response of phene states in combination is worse than that expected from the sum of their responses in isolation. For example, at low soil N levels, a phenotype with increased LRBD combined with RCA formation caused 42% reduction in shoot dry weight, compared to the expected additive effects of this phenotype, which indicates a functional antagonism (Postma and Lynch, 2011b; York et al., 2013).

A quantitative understanding of the functional dependence of one phene on the expression of other phenes and interactions among phenes and environmental factors is important for probing phenotypic diversity and breeding utility. We hypothesize that larger CCS and reduced CCFN, in combination with RCA formation, would decrease root respiration and tissue nutrient content, which would result in greater root growth, more efficient acquisition of soil N, P and K, and better root and whole plant establishment under suboptimal N, P, and K availability. We also hypothesize that the combined benefit of RCA, CCS, and CCFN is additive and is greater than the benefit of RCA alone. *OpenSimRoot*, a functional-structural plant model, was used to evaluate: (1) the utility of CCS, CCFN, and RCA under suboptimal N, P, and K availability, (2) potential synergism between CCS, CCFN, and LRBD, and (3) the benefit of CCS and CCFN under conditions of greater carbon availability as may occur with elevated atmospheric CO2 concentration.

## Results

When maize was grown under N stress in solution culture, IBM201 (genotype with reduced CCFN) showed lower N concentration in the roots, IBM30 (genotype with larger CCS) showed lower N concentration only in stems. Under sufficient P availability, both IBM201 and IBM30 showed reduced P concentration in the roots. In addition, IBM201 showed lower P concentration in leaves. Under K stress, IBM201 showed reduced K concentration in root tissues but also increased K concentration in the stems (Fig. 1).

**Figure 1.**
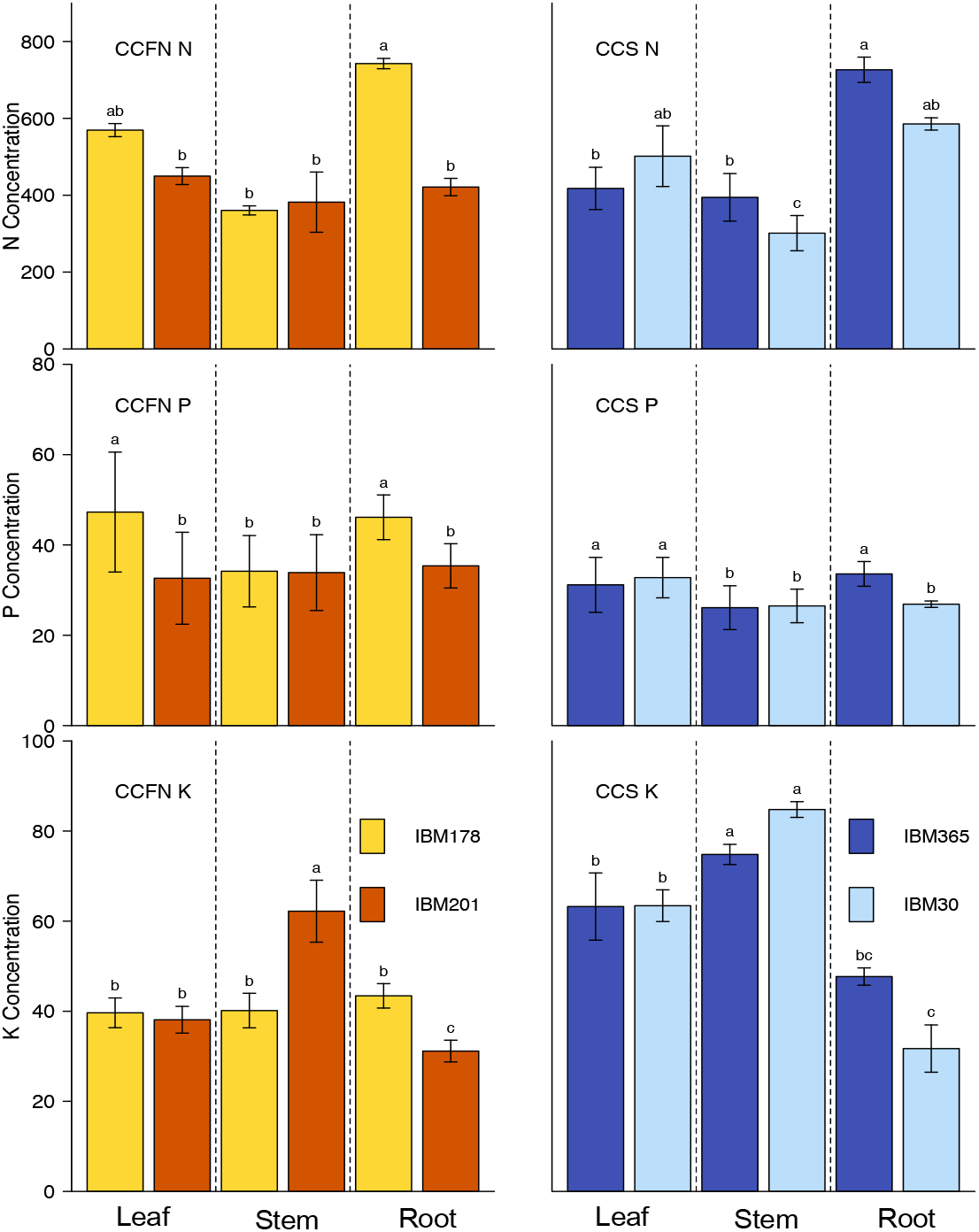
Variation in tissue nutrient concentration among maize genotypes contrasting in CCFN and CCS. Unit of tissue nutrient concentration is μmol/g dry weight. IBM178 is a many CCFN genotype, IBM201 is a reduced CCFN genotype, IBM365 is a small CCS genotype, IBM30 is a larger CCS genotype. In the nutrient solution, N concentration is 160 μmol/L, and K concentration is 60 μmol/L under N or K stress respectively. P concentration is 360umol/L. Error bars represent standard deviation of measurements from four replications.

In the simulations, increased RCA, reduced CCFN, and larger CCS had positive effects on plant growth under limiting soil nitrogen, potassium and phosphorus as simulated independently (Fig. 2, Fig. 3, Fig. 4). Plants with larger CCS and reduced CCFN had greater rooting depth and greater uptake rate of nitrate at deeper soil strata as well (Fig. 2). Improved plant growth by larger and reduced CCFN was highly dependent on the intensity of nutrient stress and the specific nutrient simulated. Generally, at intermediate deficiency (i.e., plant dry weight at 30% - 50% of an unstressed reference), RCA formation, larger CCS and reduced CCFN exhibited the greatest beneficial effect when potassium was the limiting resource, while at severe deficiency (i.e., plant dry weight at 1% - 25% of an unstressed reference), the greatest beneficial effect was found when nitrogen and phosphorus were the limiting resources. Under N and P stress the utility of all three phene states generally decreased with increasing nutrient availability, while under potassium stress they benefited the plant the most at intermediate deficiency. Reduced nutrient content in root tissue contributed more towards the total improvement in growth of phenotypes with larger CCS and reduced CCFN than did reduced respiration. The total benefits of larger CCS or reduced CCFN were greater than summing respective benefits introduced by reduced nutrient content and reduced respiration (Fig. 3), and under extreme P and K stresses, were greater than the total benefit of RCA. For example, under extremely limiting nutrient levels (21 kg/ha N, 0.05 kg/ha P), large CCS increased biomass 47% (N stress) and 56% (P stress), while the reduction in respiration contributed only 16% (N stress) and 14% (P stress), and reduced nutrient content only 18% (N stress) and 23% (P stress). Under moderate K stress (1.9 kg/ha), reduction in respiration caused by larger CCS contributed 30% and reduced nutrient content 18% towards growth benefits, while enabling both functions resulted in a 69% growth enhancements, 21% higher than the additive terms of the two functions, indicating synergism.

**Figure 2.**
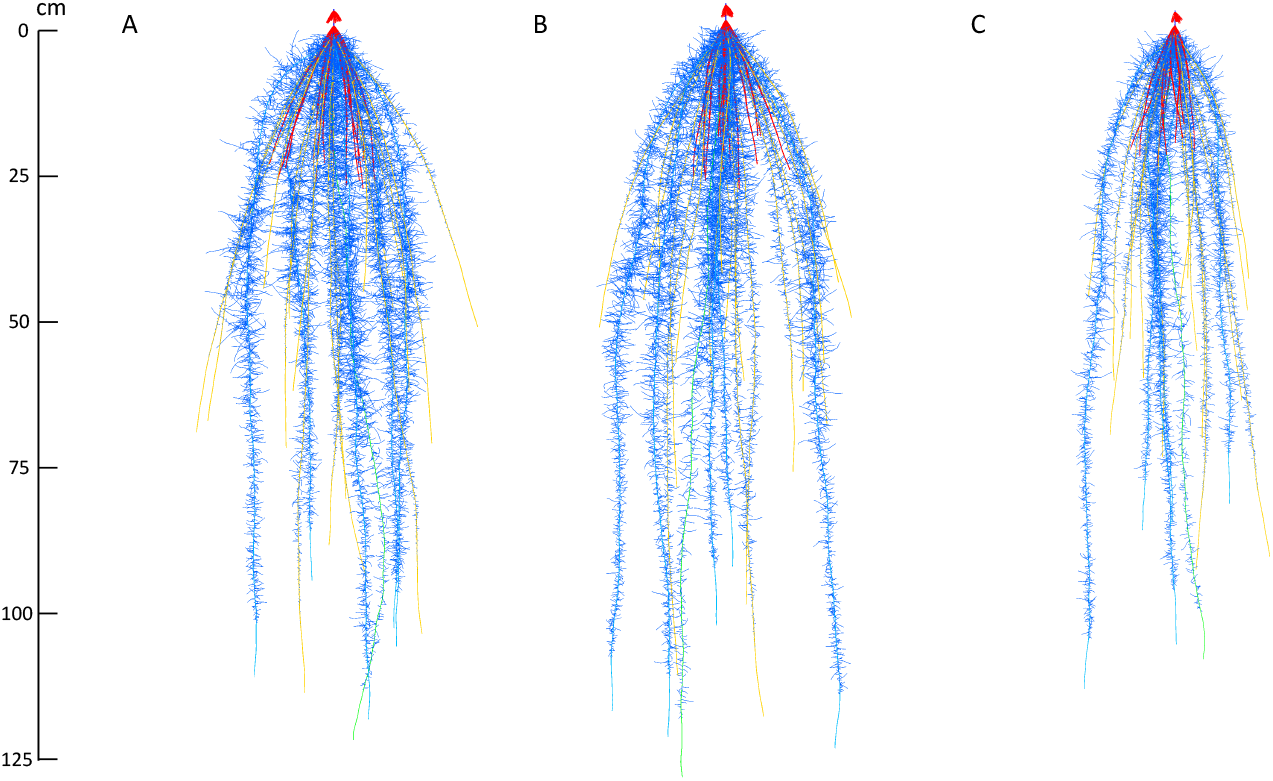
Visualized output of the simulated growth of maize root systems at 42 D.A.G. Plant growth was simulated under moderate N stress (42 kg/ha). Plant A represents a few CCFN genotype (8 cortical cell files), plant B represents a large CCS genotype (533 microns), and plant C represents a reference genotype with increased CCFN (17 cortical cell files) and reduced CCS (101 microns). The axis represents the rooting depth of the three plants in centimeters.

**Figure 3.**
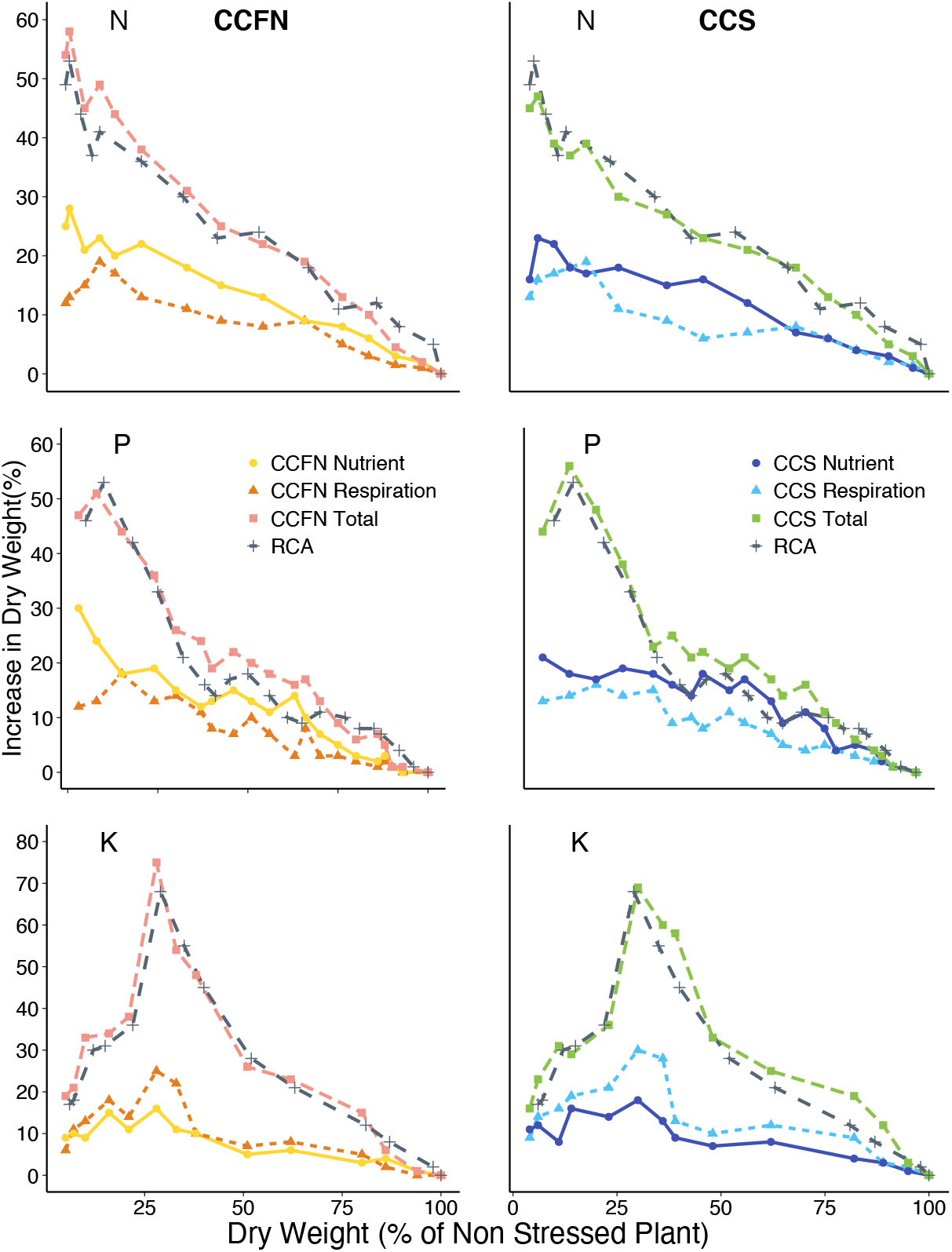
The benefits of RCA formation, larger CCS and small CCFN under suboptimal availability of N, P and K. Stresses are expressed as the relative plant dry weight at 40 days after germination compared to a non-stressed reference plant on the x axis. Benefits are expressed as increase in plant dry weight due to the presence of the phene states compared to a reference phenotype. The phenes were at the maximum beneficial level, i.e., maximum RCA formation, largest cortical cells and least cortical cell files simulated independently.

**Figure 4.**
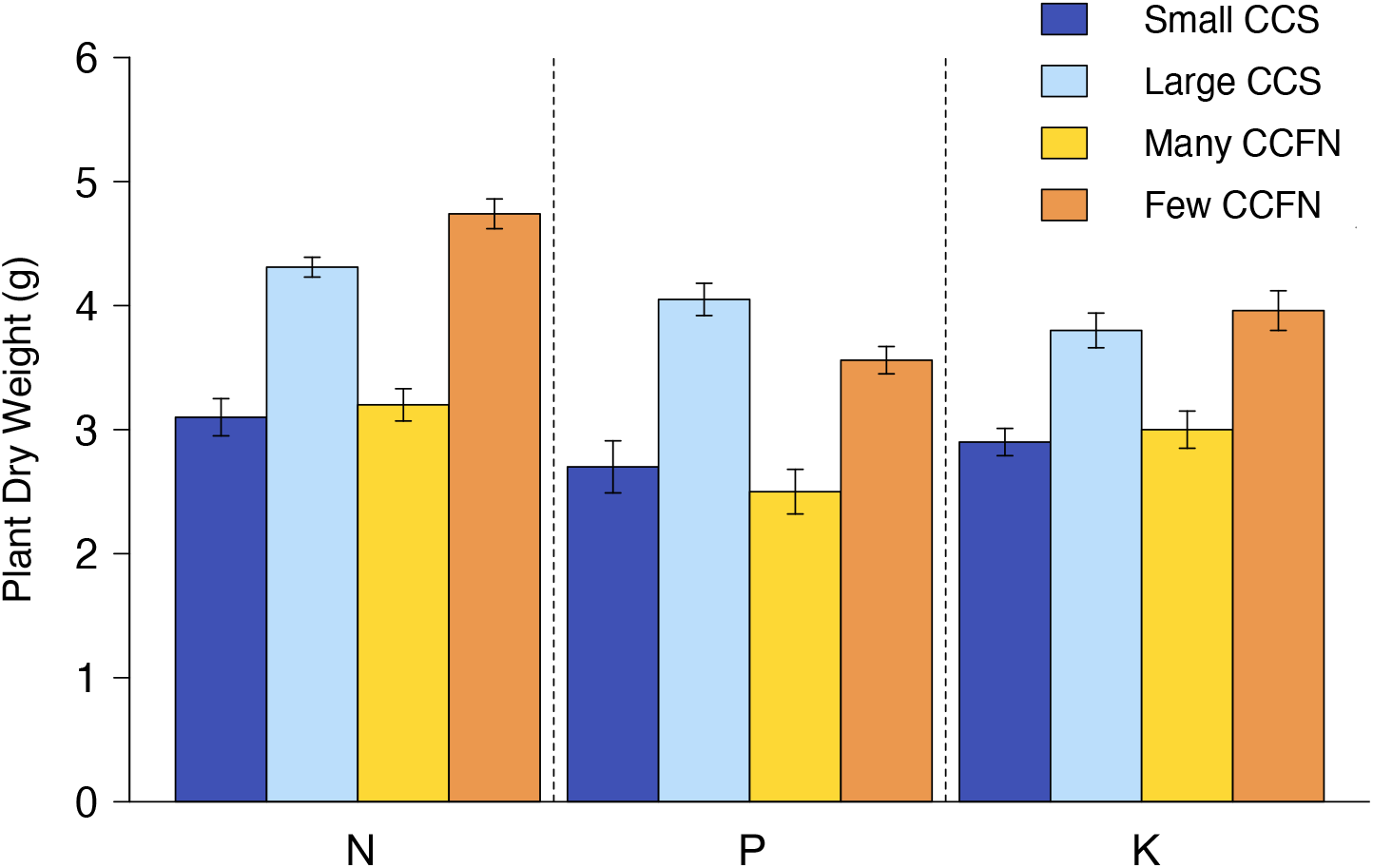
Total plant dry weight showing the utility of larger CCS (533 μm^2^) and reduced CCFN (8 cell files) vs the reference phenotype (101 μm^2^ CCS and 17 CCFN) under N, P and K stress (plant dry weight 10% of unstressed) at 40 days after germination. Error bars represent standard deviation in four repeated runs. Variation was caused by stochasticity in modeled root growth rate, root branching frequency, and root growth angle.

Predicted benefits of reducing respiration and nutrient concentration would increase in general as cell size increases and file number decreases within the range of observed variation (Fig. 5). At extremely low soil nutrient availabilities (10% of sufficient soil nitrate and potassium availabilities, 1% of sufficient soil phosphorus availability), larger CCS and reduced CCFN did not achieve the most substantial enhancement of plant growth, which were found at moderately low soil nutrient availability (20% of sufficient soil nitrate availability, 8% of sufficient soil phosphorus availability and 25% of sufficient soil potassium availability).

**Figure 5.**
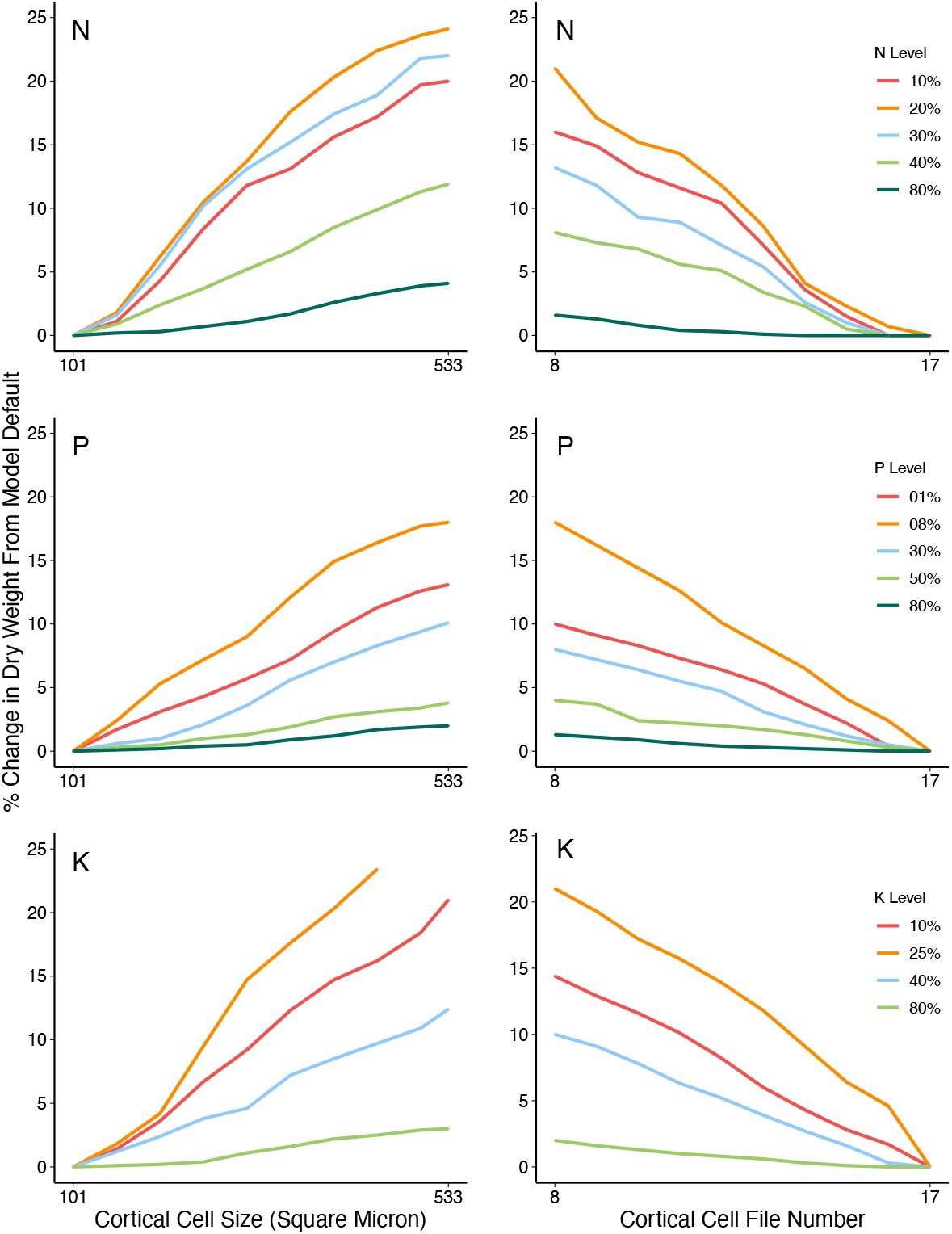
Sensitivity analysis of CCS and CCFN variation on benefits introduced by reducing respiration. Different lines correspond with percent sufficient soil nutrient availabilities as indicated. Cortical cell size and cell file number values used for simulations were based on empirical data from the literature and were within the range observed empirically. Benefits are expressed as increase in plant dry weight due to the presence of the phene states compared to the model default phenotype. N level (10% = 21 kg/ha, 20% = 42 kg/ha, 30% = 63 kg/ha, 40%= 84 kg/ha, 80% = 168 kg/ha), P level (01%= 0.05 kg/ha, 08% = 0.4 kg/ha, 30% = 1.5 kg/ha, 50% = 2.5 kg/ha, 80% = 4 kg/ha), K level (10% = 0.5 kg/ha, 25% = 1.2 kg/ha, 40% = 1.9 kg/ha, 80% = 3.8 kg/ha).

We simulated the timing and development of nitrogen and phosphorus stress in plants with only RCA formation, with both RCA formation and larger CCS, or with both RCA formation and reduced CCFN independently. Plants with either larger CCS or reduced CCFN present along with RCA formation were slightly less nitrogen and phosphorus stressed in that nutrient stress was delayed approximately 1-3 additional days by both phene states (Fig. 6). With decreases in nutrient availability, stresses developed earlier and were more severe in the phenotypes without RCA formation, or larger CCS, or reduced CCFN than the ones with these phenes states. RCA formation, CCS and CCFN could alleviate nitrogen or phosphorus stress in terms of both duration and severity of stress.

**Figure 6.**
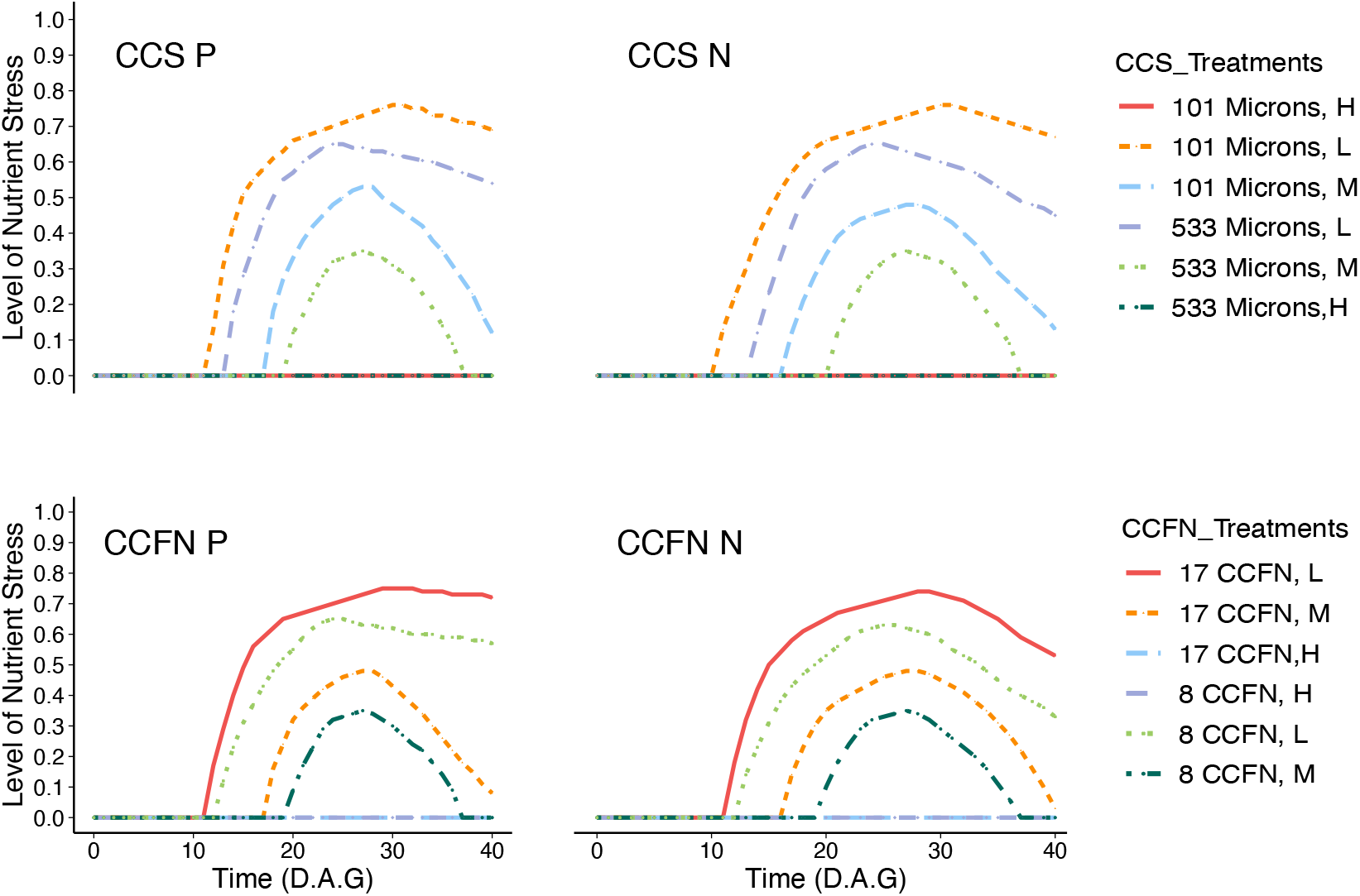
Nitrogen and phosphorus stress as affected by contrasting CCS and CCFN on a scale from 0-1. Stress is calculated as 1 – (μ – m)/(o – m), where μ is the amount of nitrate or phosphrrus being uptaken, o is the optimal nitrate or phosphorus content in the plant, and m is the minimal nitrate or phosphorus content in the plant. 0 indicates no stress, 1 indicates the most severe stress. HN = 210 kg/ha, MN = 84 kg/ha, LN = 21kg/ha, HP = 5kg/ha, MP = 2kg/ha, LP = 0.5kg/ha.

We used a high RCA phenotype with large CCS and reduced CCFN to simulate the beneficial effects of large CCS and reduced CCFN before and after RCA formation, and their interactions after RCA formation under nitrogen, phosphorus and potassium deficiency (Fig. 7). Both CCS and CCFN had initial benefits at the very beginning of growth, and both continuously increased dry weight until RCA formation, which replaced root cortical cells and cell files with large intercellular spaces. Over time, by reducing nutrient content in the root, larger CCS and reduced CCFN contributed more to growth enhancement than that of respiration reduction under N or P stress, while respiration reduction contributed more under K stress. RCA formation was at a minimal level initially, but increased substantially under suboptimal levels of all three nutrients at 15 DAG, which corresponded to the time nutrient stress was perceived due to exhausted seed reserves. After the substantial increase in RCA formation, RCA was responsible for the majority of benefits under nutrient stress. We also tested potential additive effects of RCA formation, large CCS and reduced CCFN, since the majority of benefits of RCA do not overlap with those of large CCS and reduced CCFN over time during growth. Benefits of both large CCS and reduced CCFN after RCA formation at 15 DAG were reduced. After 15 DAG, the majority of benefits were contributed by RCA. In this case, the combination of all three phene states at their most carbon-efficient level, i.e., the observed level that showed greatest reduction in the carbon cost of root maintenance, achieved growth benefits up to 135%, 132% or 161% under low nitrate, phosphorus or potassium availabilities.

**Figure 7.**
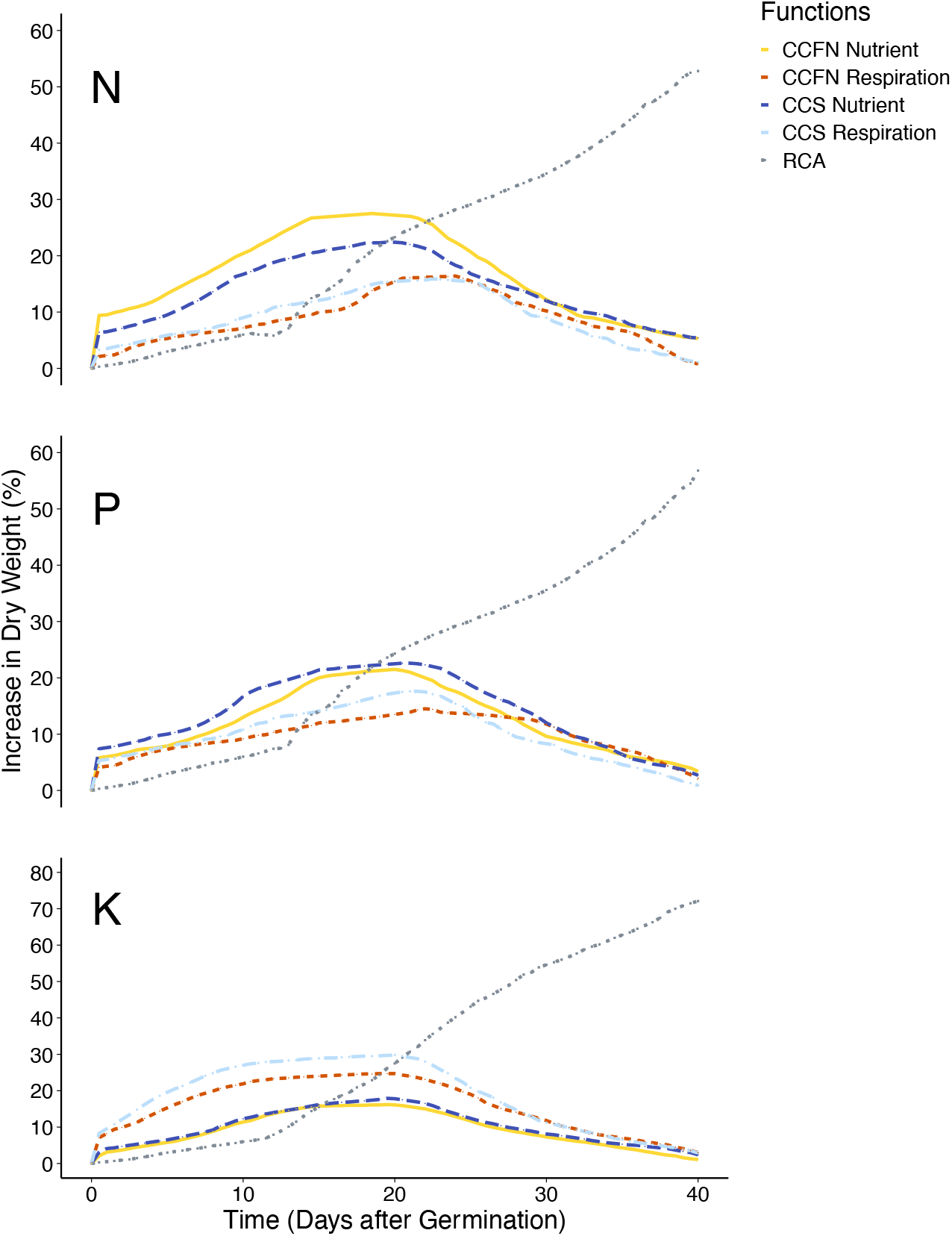
Relative benefit of having RCA formation, larger CCS and small CCFN simultaneously in a simulated plant over time at 42 kg/ha of soil nitrate, 0.5 kg/ha of soil phosphorus and 1.5kg/ha of soil potassium availability. Different lines correspond to relative benefits for respiration and nutrient content, similarly described in figure 2. The gray line indicates the hypothetical additive benefit when RCA formation achieves the optimal growth enhancement

We simulated the utility of CCS and CCFN under nitrogen and phosphorus deficiency with varied lateral root branching density (LRBD) in phenotypes with either the largest root cortical cells, or fewest root cortical cell files. Both large CCS and reduced CCFN achieved greater growth enhancements in phenotypes with half the reference LRBD under low soil nitrate availability. The contrary was evident for phenotypes grown under both low and medium soil phosphorus availability, where greater benefits were observed when plants with doubled LRBD compared to the reference phenotype. In soils with intermediate nitrate availability, phenotypes with normal LRBD had the greatest benefit compared to half or doubled LRBD (Fig. 8).

**Figure 8.**
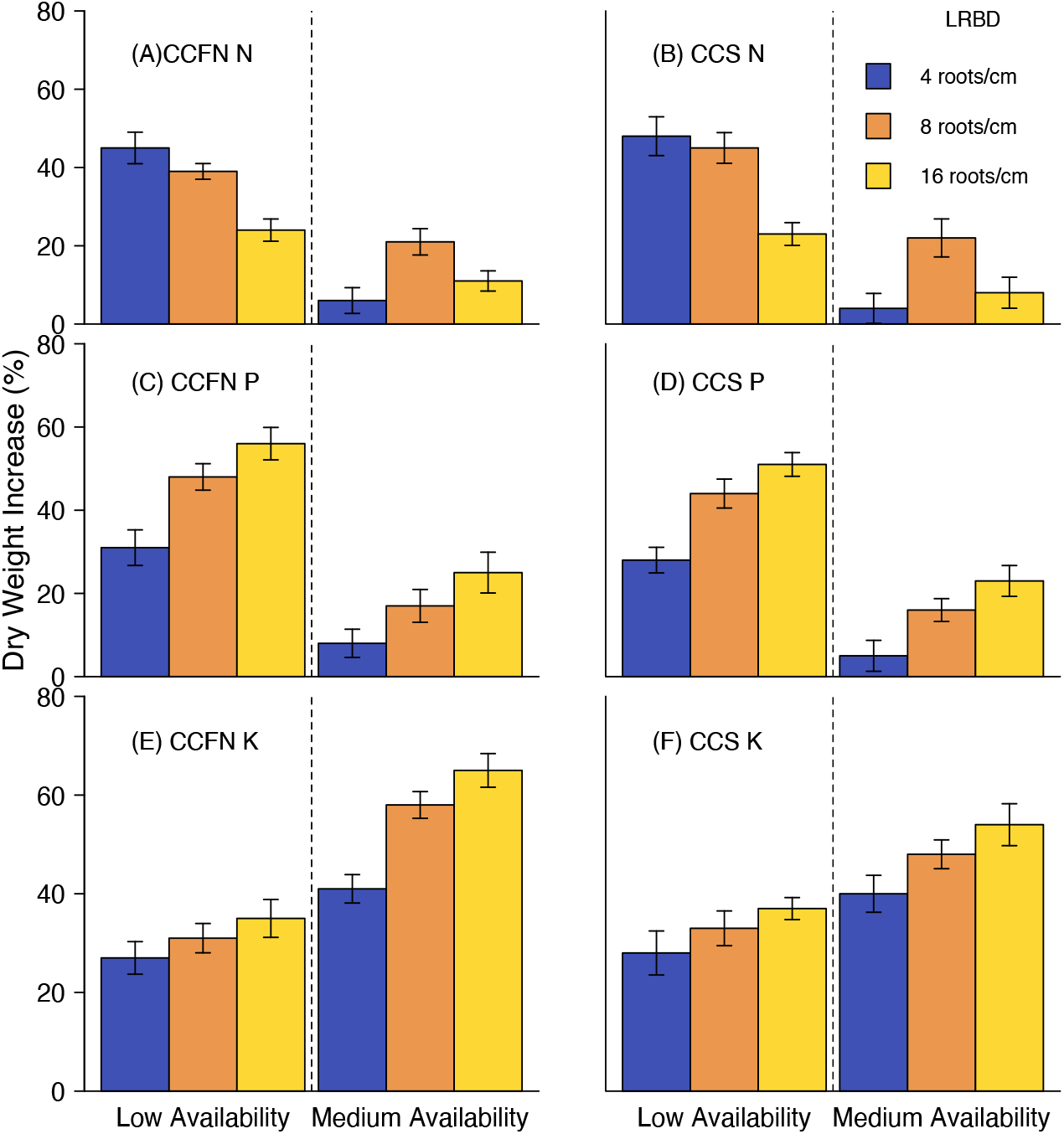
Interactions between larger CCS and Lateral Root Branching Density (LRBD) (A, C and E), and reduced CCFN and LRBD (B, D, and F) under low or medium soil nitrogen, phosphorus or potassium availability. CCS and CCFN used in this scenario are both at the least level of metabolic carbon demand (largest cell size, reduced cell files). Three levels of Lateral Root Branching Density (4, 8, and 16 roots/cm) are shown, the range of which was based on Trachsel et al. (2010), as used by Postma and Lynch (2011). Low and medium nitrate levels are 21 kg/ha and 84 kg/ha respectively. Low and medium phosphorus levels are 0.5 kg/ha and 2 kg/ha respectively. Low and medium potassium levels are 0.5 kg/ha and 1.9 kg/ha respectively.

With greater carbon availability, simulated by increasing light utilization efficiency in the canopy module, the benefits of large CCS under low soil nitrogen or phosphorus availabilitydeclined (Fig. 9).

**Figure 9.**
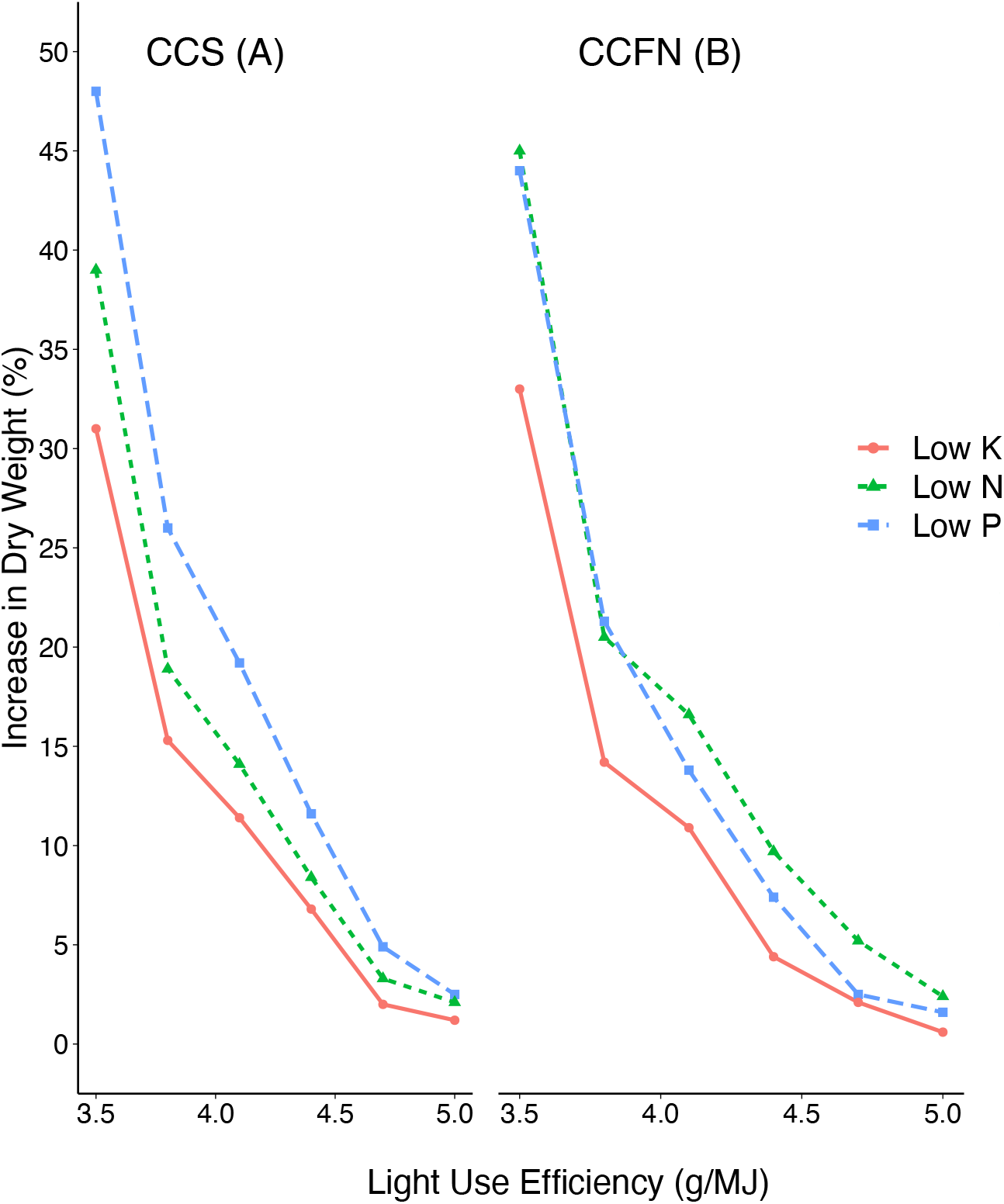
Benefits of CCS and CCFN on plants grown under nitrogen or phosphorus deficiencies as affected by elevated atmospheric CO2 concentration, as simulated by increased canopy light use efficiency. Soil phosphorus level was 0.4kg/ha, nitrate level 21kg/ha, and potassium level 1kg/ha.

## Discussion

Our results align with previous findings that RCA formation, which reduces the metabolic cost of soil exploration in terms of nutrient and C investment, improves plant growth under conditions of suboptimal availability of N, P and K (Postma and Lynch, 2011; Saengwilai et al., 2014; Galindo-Castañeda et al., 2019), and support the hypothesis that larger CCS and reduced CCFN increase soil nutrient acquisition by reducing root metabolic costs. The combined benefits of RCA formation, larger CCS and reduced CCFN for growth are greater than the benefit of RCA formation alone. Larger CCS and reduced CCFN reduce the metabolic cost of soil exploration under drought stress (Chimungu et al., 2014a, b), and are predicted by our results to alleviate nitrogen, phosphorus and potassium stress as well. No literature has reported CCS or CCFN alleviating potassium stress. However, given that both larger CCS and reduced CCFN reduce root respiration (Chimungu et al., 2014a, b), we believe that they may have utility under potassium stress. These results indicate that all three phenes may have substantial utility on infertile soils, suggesting that cultivars with high RCA formation, large cortical cells and reduced cortical cell files would have reduced fertilizer requirements in intensive agriculture and would yield better in low-input systems. Our results focus on maize but we propose that they should be generally applicable to other grass species, which like most monocots lack secondary growth and so have a persistent cortex.

Typically, multiple edaphic stresses occur simultaneously (St. Clair and Lynch, 2010), although it is difficult to reflect concurrent stresses in the model because of potential interactions among plant stress responses (Dathe et al., 2013). In such environments, tradeoffs between nutrient acquisition strategies for specific nutrients, and between other plant physiological functions, are challenging (Hu et al., 2014; Lynch and St. Clair, 2004; Rubio et al., 2003). For example, the strategy of enhancing topsoil foraging proved to be critical for adaptations to low phosphorus soil in common bean (Lynch and Brown, 2001) and maize (Zhu et al., 2005); the “Steep, Cheap and Deep” ideotype proposes that root phenotypes capable of rapid exploration of deep soil strata would optimize soil nitrate and water capture in maize (Lynch, 2013). However, given the limited amount of carbon and nutrients available for root maintenance, extreme inclinations towards one strategy may be detrimental for the other, and optimization of resource allocation between deep soil exploration and topsoil exploration is critical. For example, the optimal lateral root branching density in maize is dependent upon nitrate and phosphorus availability (Postma et al., 2014). Sparse but long LRBD is optimal for nitrate uptake, while dense but short LRBD is optimal for phosphorus uptake. In another study (Dathe et al., 2016), axial root growth angle exhibit significant effects on nitrogen acquisition in maize, where extreme phenotypes have narrow intervals of optimal performances – extremely shallow root systems only increase N acquisition under reduced precipitation, while dimorphic phenotypes that combined shallow seminal roots with deep crown roots performed well in all environments. High LRBD introduces strong competition among roots for nitrate capture, therefore decreasing nitrate uptake since the carbon budget of the whole plant does not grant greater root length. Carbon budget and root competition does not impact phosphorus uptake as significantly as nitrate uptake, therefore increasing root length by increasing LRBD is optimal for phosphorus uptake. In reality, most genotypes have a balanced LRBD to meet the demand of both nitrate and phosphorus acquisition. Maize genotypes with high RCA formation could also inhibit radial phosphorus transport due to the reduction of living tissue (Hu et al., 2014).

The utilities of RCA, larger CCS and reduced CCFN were greater in plants that were experiencing moderate potassium stress (25% of potential growth) than in plants under severe potassium stress (less than 10% of potential growth). Similarly, RCA, larger CCS and reduced CCFN did not achieve optimal growth enhancement under extremely low soil nitrate and phosphorus availability (1% to 8% of potential growth under N stress, 1% to 5% of potential growth under P stress), but showed greater benefits under less severe stress. This decline is caused by reduction in the utility of the respiration reduction function. With extremely limited nutrient availability, root respiration per ion absorbed increases, inhibiting root and shoot growth to compensate for greater respiration. The reduction in respiration is more important in potassium deficient plants than in nitrogen deficient or phosphorus deficient plants, as carbon is relatively more limiting in potassium stressed plants, which differs from the cases of nitrogen and phosphorus (Postma and Lynch, 2011a). Substantial reductions in photosynthetic assimilation caused by nutrient deficiency impose carbon limitations under N, P or K stress. However, in potassium stressed plants, an adaptive response in carbon partitioning between roots and shoots is absent, which is present in nitrogen or phosphorus stressed plants, that can allocate more carbon to root growth.

The model predicts a decline in the benefit of larger CCS and reduced CCFN at extremely low nutrient availability. If we consider the amount of nutrient required for the construction of a root segment as the cost, then the cost is greater in low nutrient soils than in fertile soils (Postma and Lynch, 2011b). Nutrient uptake under severe deficiency could be limited by both carbon and the deficient nutrient, while the cost of root tissue construction and respiration remain high, causing an increase in the cost-benefit ratio for nutrient uptake (Nielsen et al., 2001). In *OpenSimRoot*, the benefit of larger CCS and reduced CCFN is dependent on the cost – benefit ratio of root segments. Under extremely scarce nutrient availability, the decline in benefits is caused by the high cost – benefit ratio of root growth. Therefore, if nutrient availabilities fall below a threshold where the stressed plant does not have sufficient nutrient stores for tissue construction, causing extra allocation of carbon to the root system, thus deteriorating the photosynthetic and nutritional status of the plant, then the utility of RCA formation, CCS, CCFN, or other phenes that reduce metabolic costs, would decrease. In an extreme theoretical environment where all nutrients in soil are depleted, the utility of these phenes becomes nil.

While the potential for RCA formation is genetically controlled, RCA formation is highly responsive to edaphic stress (Fan et al., 2003). Substantial variation in CCS and CCFN, however, were observed among RILs in empirical studies, but were not as plastic to edaphic stress as RCA formation (Chimungu et al., 2014a, b), and could be beneficial starting at the very beginning of growth under edaphic stress. In our simulation, both large CCS and reduced CCFN exhibited benefits before the formation of RCA in response to nutrient limitation. As cortical cells and cell files were replaced by RCA formation, we expect the majority of benefits from these three phenes do not overlap over time. Therefore, we predict a simplification of the additive effects among the three phenes to be present under N, P or K stress, and observed increased benefits due to the combination of all three phene states compared to that of RCA formation alone. In reality, however, we expect a more complicated interaction between RCA formation, CCS and CCFN. RCA formation, as a response to nutrient stress, manifested 12-13 days after germination (Postma and Lynch, 2011). RCA formation in mid-root and apical regions were less than that of basal regions of a root (Fan et al., 2003). The variation in the spatial and temporal distribution of RCA at both the single root scale and the root system scale is dynamic (Burton et al., 2013), and such variation can cause changes in nitrogen uptake kinetics (York et al., 2016). Therefore, in reality, the interaction between RCA formation, CCS and CCFN could not be represented by a simplified expectation of additive effects.

Plants have developed multiple alternate strategies to increase nutrient uptake under severe nutrient stress, such as root hair formation, root exudation, and mycorrhizal colonization. Root hair formation has a relatively low cost – benefit ratio, but can increase phosphorus uptake significantly (Bates and Lynch, 2001; Nielsen et al., 1994, 2001; Zhu and Lynch, 2004; Miguel et al., 2015). Mycorrhizal colonization increased phosphorus efficiency significantly at low phosphorus availability compared to non-colonized plants (Ning and Cumming, 2001), but did not increase plant dry weight significantly due to the increased maintenance and growth respiration of the fungal tissue (Nielsen et al., 2001), indicating that root carbon costs are a major limitation to plant growth under phosphorus stress. Additionally,when the carbon cost of root growth is removed, simulated plant growth increased under P stress (Postma and Lynch, 2011). Our results support the general hypothesis that the metabolic costs of soil exploration in terms of the carbon and nutrient investment in root tissue over time becomes increasingly important as the availability of crituical soil resources declines.

Chimungu (2014) reported, in both drought-stressed and non-stressed plants, significantly thicker roots when larger cortical cells and more cortical cell files were present. In addition, Burton (2010) observed greater RCA formation in thicker root classes in non-stressed plants. The ability of roots to penetrate compacted soil and root depth are correlated to both root anatomical phenes such as RCA, CCS and CCFN, as well as root diameter, where deeper-rooting plants in compacted soil showed reduced CCFN and increased RCA formation. Additionally, root thickening in the form of root cortical area expansion were closely related to soil mechanical impedance in some genotypes (Chimungu et al., 2015b; Vanhees et al., 2020). Smaller outer band cortical cells could reduce the risk of root collapse when encountering increased mechanical impedance. RCA formation is also negatively correlated with root bending strength, while smaller distal root cortical cells, more cortical cells, and more CCFN increase the strength of root and reduce root collapsing during penetration of soil (Whiteley et al., 1982; Clark et al., 2003; Jin et al., 2013;) since they contribute to the construction of a thicker root. Root cortical cell diameter is a pivotal and heritable trait in determining the carbon cost of penetrating compacted soil, where large cell diameter correlated with reduced carbon cost of root growth, especially under high soil mechanical impedance; the plasticity of this trait allowed the enlargement of root cortical cells was a common response when roots encounter compacted soil (Colombi et al., 2019). These observations from empirical studies suggest that, although RCA formation, larger CCS and reduced CCFN reduce the metabolic cost of soil exploration, they may affect root penetration of hard soil domains, which could affect their ability to acquire resources located in deeper soil profiles.

An important merit of simulation modeling is the ability to test hypotheses and probe scenarios that are inaccessible to empirical studies. Simulation modeling makes it possible to isolate and test the objects of study, in this case, larger CCS and reduced CCFN, from interactions with many other biotic and abiotic factors, which is difficult to avoid in empirical studies (Dunbabin et al., 2013; Postma et al., 2014). It would be infeasible to test separately how larger CCS and reduced CCFN reduce respiration and nutrient content in an empirical study, while in modeling, different functions of specific phenes could be isolated and examined without being confounded by other functions, which was critical to several previous studies (Postma and Lynch, 2011a; 2011b). In other cases, modeling also allows us to study phenes that are otherwise difficult to manipulate in real plants, such as changing nutrient uptake kinetics (York et al., 2016), or examining root competition in time and space in the ‘three sisters’ polyculture (Postma et al., 2014) where empirical measurement is impractical. *In silico* approach allows the flexibility of conducting thousands of simulations in factorial designs (in this study, over 3,300 runs) which would be difficult to conduct empirically. Although such models are designed with assumptions and simplifications of the actual scenarios or mechanisms they simulate, and often (as in the present case) rely on empirical data as input parameters, it does not nullify the value of models as a useful research tool to provide a preliminary insight into root anatomy, architecture, physiological processes and interactions with other factors of interest, and a compliment to field studies even when empirical data are present. In our case, *OpenSimRoot* is capable of simulating a comprehensive range of phenes and phenotypes in specific environments that can be customized. Because of its heuristic nature, *OpenSimRoot* focuses on the validity of simulating physiological processes, rather than the alignment with empirical studies that predictive models emphasize.

Our results suggest potential areas where structural-functional plant models could be improved. The duration of simulation could be parameterized for longer periods, which could potentially enable models to simulate the full life cycle of plants, and demonstrate the dynamics of physiological processes. Interactions and dynamics among root phene states deserve more attention. Some other parameters, such as soil hardness, microbial associations, interplant competition (Postma and Lynch, 2011), and interspecific interactions in cropping systems (Postma and Lynch, 2012) are important for understanding ecosystem functioning on a greater scale, and may have consequences for the utility of root anatomical phenes such as RCA, CCS and CCFN. The phene aggregate of reduced CCFN and larger CCS, along with RCA formation, is defined as reduced living cortical area (LCA). Plants with reduced LCA had decreased root segment respiration, reduced P concentration in root tissues, and greater rooting depth, indicating lower carbon cost of root growth, which resulted in increased biomass and resource capture under P stress (Chimungu et al., 2014b; Galindo-Castañeda et al., 2019).

## Conclusions

Quantitative evidence that larger CCS and reduced CCFN are adaptive phene states for multiple nutrient stresses are presented. The utilities of larger CCS and reduced CCFN in soils with suboptimal nitrogen, phosphorus and potassium availability are dependent on nutrient availability, phene functions, and interactions among phenes. We propose that larger CCS and reduced CCFN are complementary to RCA formation in terms of growth enhancements as the majority of benefits of these three phene states do not overlap in time. We expect tradeoffs for RCA formation, larger CCS and reduced CCFN to be present, as all three phene states are related to soil penetration and root proliferation. This aspect merits further investigation. Functional-structural plant models like *OpenSimRoot* can be used to simulate variations in these anatomical phenes and thereby evaluate their utilities under multiple edaphic stresses, and have the potential to provide a holistic understanding of the roles of root phenotypes for plant fitness. These results indicate that large CCS and reduced CCFN merit investigation as breeding targets for maize and possibly other cereal crops, since the development of crop cultivars with improved soil resource acquisition remains a critical strategy for improving the sustainability of intensive agriculture and for improving the productivity of low-input agroecosystems.

## Materials and Methods

We used *OpenSimRoot* (Lynch et al., 1997), a functional-structural plant model with focus on root architecture and soil resource acquisition, to simulate the formation of RCA, variation in CCS and CCFN, and their physiological utility in maize growing with varied nitrate, phosphorus, or potassium availability in the soil. We also evaluated potential additive effects among RCA, CCS, and CCFN. In addition, we conducted a pair of two factor solution culture experiments to examine the variation in tissue N, P and K concentration in genotypes with contrast in CCS or CCFN.

### Solution culture study

Four maize genotypes (IBM population IBM178, IBM201, IBM365, IBM30) were used in the solution culture study. Genotypes were selected for contrasting CCS (IBM365 and IBM30), and CCFN (IBM178 and IBM201) based on preliminary screening.

Genotypes were planted in four replications in total under both high and low N or K availabilities in solution culture, with sufficient P availability across all treatments in a greenhouse at the Penn State University campus located at University Park, PA, USA (40.8148° N, 77.8653° W), with two replications planted on June 4^th^, and two more replications planted on June 13^th^, 2017. Eight 100-liter non-transparent plexi glass solution culture tanks were used, within each tank, two replications of the four genotypes were planted. Nutrient solution was based on and modified upon the Hoagland solution (Johnson et al., 1957). N concentration in N stress treatments was 160umol/L, and K concentration in K stress treatments was 60 umol/L, P concentration was 2mmol/L.

Plants were grown for 24 days after transplanting, or 32 days after germination in growth chamber to avoid RCA formation confounding the effect of larger CCS and reduced CCFN. Upon harvest, 10 cm long root segments from base and tip of the second and the third whorl nodal roots were collected to conduct anatomical analysis and respiration measurements. The rest of the root system, along with leaves and stems, were separated and dried in oven at 60 °C to measure root and shoot dry weight. Tissues were then ground and sent to the Agricultural Analytical Services Laboratory at the Penn State University for P and K content analysis. N content analysis was conducted with a 2400 CHNS/O Series II element analyzer (PerkinElmer). Nutrient content data was used to parameterize *OpenSimRoot* to include how tissue nutrient content was reduced by larger CCS and reduced CCFN in the simulations.

### Model description

*OpenSimRoot* simulates the three-dimensional root architecture and soil resource acquisition of a root system over time. The root system is described as distinct root classes represented by a growing number of root nodes and segments as the root system develops (Lynch et al. 1997). Root growth is based on a carbon source-sink model, where the carbon partition protocol has been described by Postma and Lynch (2011a). Shoot growth and photosynthesis is simulated using LINTUL (Spitters and Schapendonk, 1990). Nutrient uptake is simulated for each root segment in comparison with the optimal (o) and minimal (m) nutrient requirements of the plant. Nutrient deficiency, or stress factor, is defined as when nutrient uptake falls below the optimal nutrient requirement. The stress factor influences shoot development and photosynthetic efficiency depending on the nutrient simulated. We used the Barber-Cushman model (Itoh and Barber, 1983; Postma and Lynch, 2011a) to simulate phosphorus uptake, and linked *OpenSimRoot* to the three-dimensional hydrological model SWMS3D (Simunek et al., 1995) to simulate nitrate and potassium uptake. The Barber-Cushman model is considered to be inadequate for nitrate and potassium uptake as these nutrients are relatively mobile (Postma and Lynch, 2011b), and the Barber-Cushman model does not simulate leaching and ignores root competition in three dimensions. The SWMS3D model is not ideal for simulating the phosphorus depletion zones at root surface (Postma and Lynch, 2011b), as computational demands required by the resulting substantial number of finite element (FEM) nodes (Hardelauf et al., 2007) are considerable, and the phosphorus depletion zone would be artificially enlarged in the SWMS3D model.

Variation in RCA formation, CCS and CCFN in maize are simulated for each root segment with empirical parameters retrieved from Burton (2010), and Chimungu (2014a, b). The percentage RCA for different root classes is well described by Fan et al. (2003). We simplified CCS and CCFN simulation by assuming they are uniformly distributed across root classes. The addition of CCS and CCFN were implemented as new model input files, no specific modification were made to the computational codes of OSR to accommodate this addition. RCA formation is allowed to combine nutrient remobilization and respiration reduction and is based on regression between the amount of RCA and nutrient content and root respiration of empirical measurements by Fan et al. (2003). *OpenSimRoot* does not explicitly represent root anatomy, so CCS and CCFN variation is represented by reducing modeled root respiration and tissue nutrient content.

### Effects of nutrient stress on growth

In *OpenSimRoot*, the nutrient stress factor module is implemented to affect the potential leaf area expansion rate and light use efficiency (LUE) independently as in the LINTUL model. The nutrient stress factor functions as a growth regulator between root and shoot growth. The nutrient stress factor negatively impacts light use efficiency and resulted in reduced carbon available for plant growth. Reduction in the potential leaf area expansion rate caused by the stress factor resulted in reduced sink strength of the shoot, and consequently greater allocation of carbon to root growth. Nutrient-specific stress response was used to determine the effect of internal nutrient concentrations (nitrogen, potassium and phosphate) on the two parameters. In this study, potassium stress strongly reduces LUE (Zhao et al., 2001) but does not affect the potential leaf area expansion rate (Cakmak et al., 1994). Suboptimal phosphate strongly reduces the potential leaf area expansion rate but is trivial in affecting LUE (Lynch et al., 1991). Inorganic nitrogen strongly affects both parameters (Uhart and Andrade, 1995).

### Distribution of RCA formation, CCS and CCFN within the root system

We assumed that RCA formation starts behind the elongation zone of a root and develops over time until reaching a maximum. Therefore, the greatest amount of RCA formation can be found close to the base of a root, which aligns with Fan et al. (2003) but disagrees with Bouranis et al. (2006), Lenochová et al. (2009), and Burton (2010). RCA formation is reduced in the first 5 cm of the root (Bouranis et al., 2006), which is a small part of the total root length and we expect the effect on total RCA formation to be small. We used the maximum amount of RCA formation in the literature, which is 39% of root cross-section area at 20 days after germination (Fan et al., 2003) in the model.

Variation in CCS and CCFN was observed in the mid cortical band of roots by Chimungu et al. (2014a, b). In reality, the spatial distribution of CCS and CCFN are not uniform in either the area cross-sectioned, or across different root classes. To demonstrate the effect of observed respiration reduction of these phenes, we assumed that CCS and CCFN variation are uniform regardless of root class and location in the area cross-sectioned. Parameterization of these phenes are based on the genotypic variation described by Chimungu et al. (2014a, b). CCS varies between 101 μm^2^ and 533 μm^2^, and CCFN varies between 8 and 17 in maize.

### Interactions between RCA formation, LRBD, CCS and CCFN

Living cortical area (LCA; Jaramillo et al., 2013) is proposed as a good predictor of root respiration, and a critical determinant of root metabolic cost, which involves the phenes in this study. Interactions between LCA components requires further demonstration. We simulated the extremes of variation in RCA formation, where RCA takes up between 0% to 39% of the root cross sectional area, CCS and CCFN to probe additive effects under low nitrogen and phosphorus availability. Significant genetic variation exists in lateral root branching density (LRBD) (Trachsel et al., 2011). We varied the LRBD parameter to the extremes reported in this study, between 4 to 16 lateral roots/cm on axial roots, to examine if potential synergism between LRBD, CCS and CCFN under low soil nitrogen and phosphorus availability to further test the utility of CCS and CCFN in an integrated genotype.

### System description, parameterization, and runs

We simulated growth of 40 days after germination of a single maize plant, which represents a uniform monoculture plant community with a between-row spacing of 60 cm and a within row spacing of 26 cm. Aboveground competition was simulated by a shading function (Postma and Lynch, 2011). Parameterization was based on input parameters used in previous simulation studies with *OpenSimRoot* (Postma and Lynch, 2011a; 2011b). From empirical measurements from Chimungu *et al*. (2014a, b), we parameterized how larger CCS and reduced CCFN reduced root respiration. We parameterized how tissue nutrient content varied between contrasting phenotypes by conducting a solution culture study (see above). All simulations were performed on the Penn State supercomputing clusters aci-b, with the following variables: (1) CCS and CCFN; (2) the functions of RCA formation; (3) lateral root branching density with CCS and CCFN held constant; (4) atmospheric CO2 pressure between ambient values of 400ppm, and up to four-fold (1600 ppm); (5) the availability of nitrate, phosphorus and potassium in the soil, from low to sufficient; and (6) “max and min RCA”, “max and min CCS” and “max and min CCFN” reference genotypes. To account for stochasticity in growth rates and root branching frequencies, each scenario was simulated with four replications each with *OpenSimRoot’s* random number generator initialized to different values, with the graph showing the mean value. The variation of phenes in this study were based on empirical studies to avoid extrapolation towards unrealistic conditions. Appendix A contains a summary of the model parameterizations.

### Statistical analysis

Empirical data from the solution culture study were analyzed by paired Student’s *t* tests in R 3.4.1 (R Core Team, 2017). We did not conduct significance test on the simulation results, as such tests were not reliable in simulation studies, as the ease of replication in computer simulations allows for any effect size to be found significant if there are enough replicates. Biological significance, rather than the statistical significance, should be the main focus of simulation experiments (White et al., 2014).

## Abbreviations

(CCS): Cortical cell size
(CCFN): Cortical cell file number
(RCA): Root cortical aerenchyma
(SCD): Steep, Cheap and Deep
(LRBD): Lateral root branching density
(RHL): Root hair length
(BRGA): Basal root growth angle

## Appendix OpenSimRoot Parameterization

OpenSimRoot uses a hierarchical xml formatted input file which is graphically presented below. The hierarchy places the parameters in a context. For example, the parameter ‘specific leaf area’ belongs to the shoot of a specific genotype. In OpenSimRoot parameters can be a single value, a value drawn from a distribution, or the result of an interpolation table. We have tried to base all our parameters on our own measurements or data from the literature. We have indicated the sources behind the parameters. Note that in many cases, we used more than one source and in some cases we had to convert the measurements using assumptions. A common assumption we made is that the value was equal for all root classes and or for all genotypes.

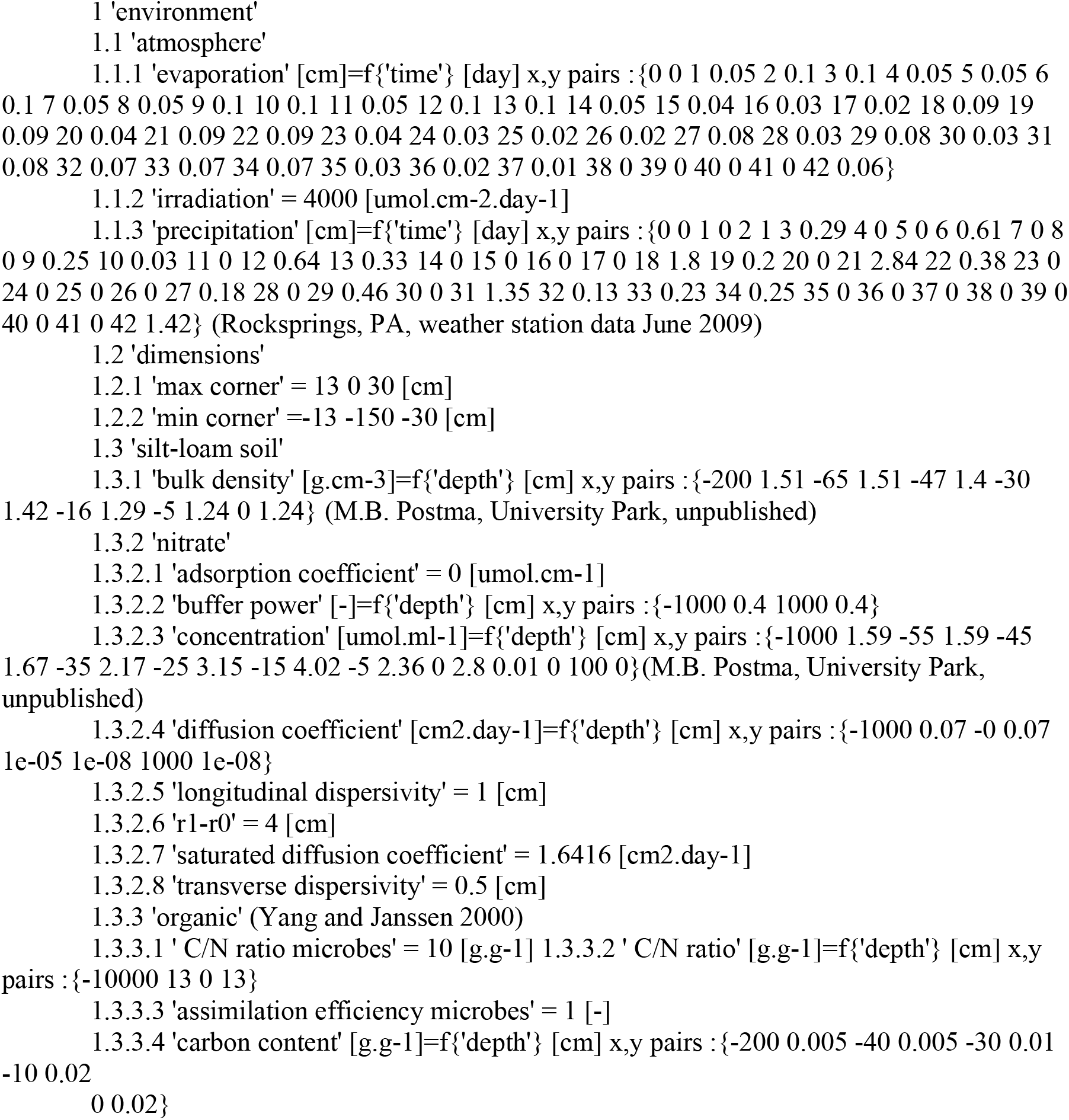
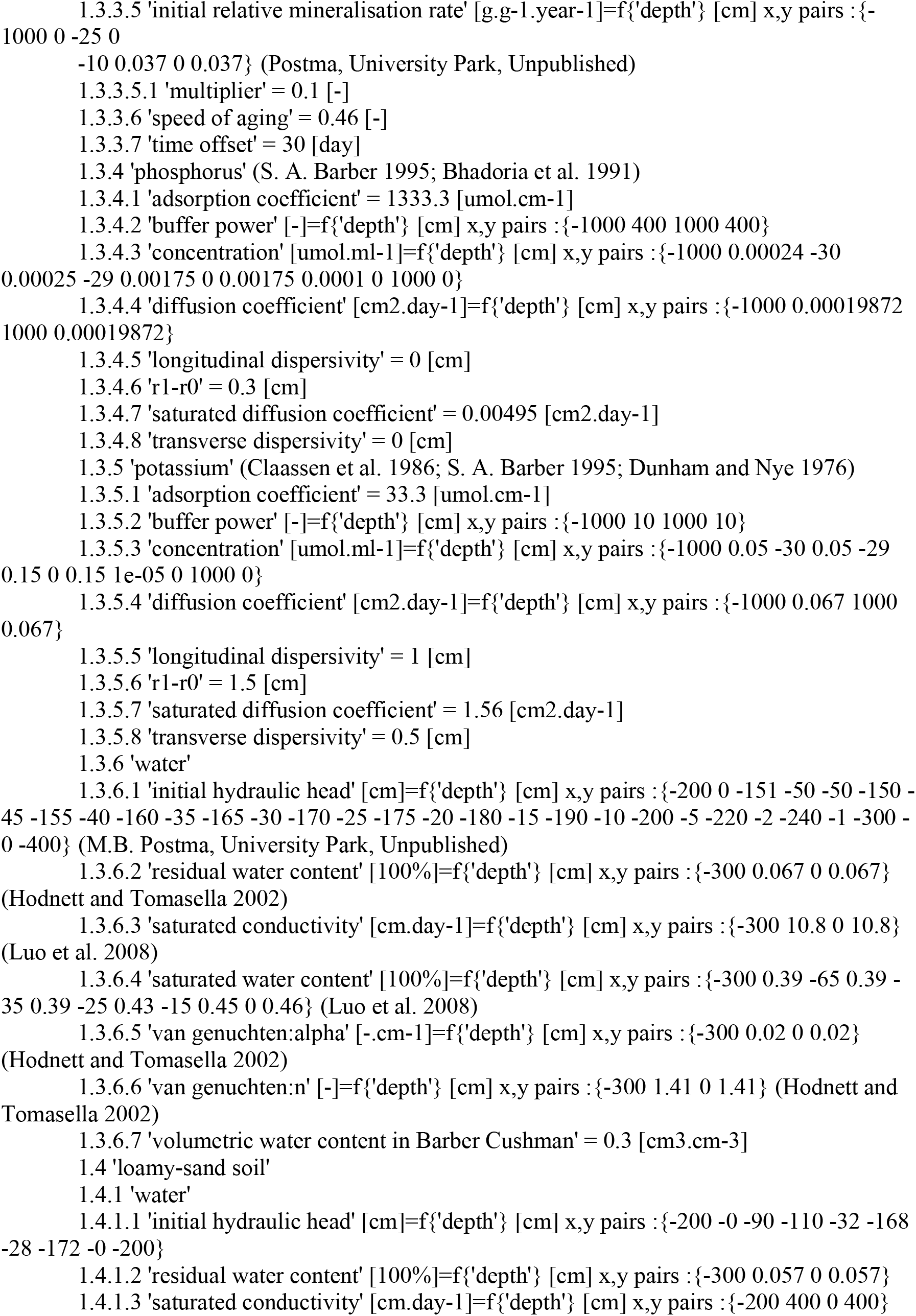
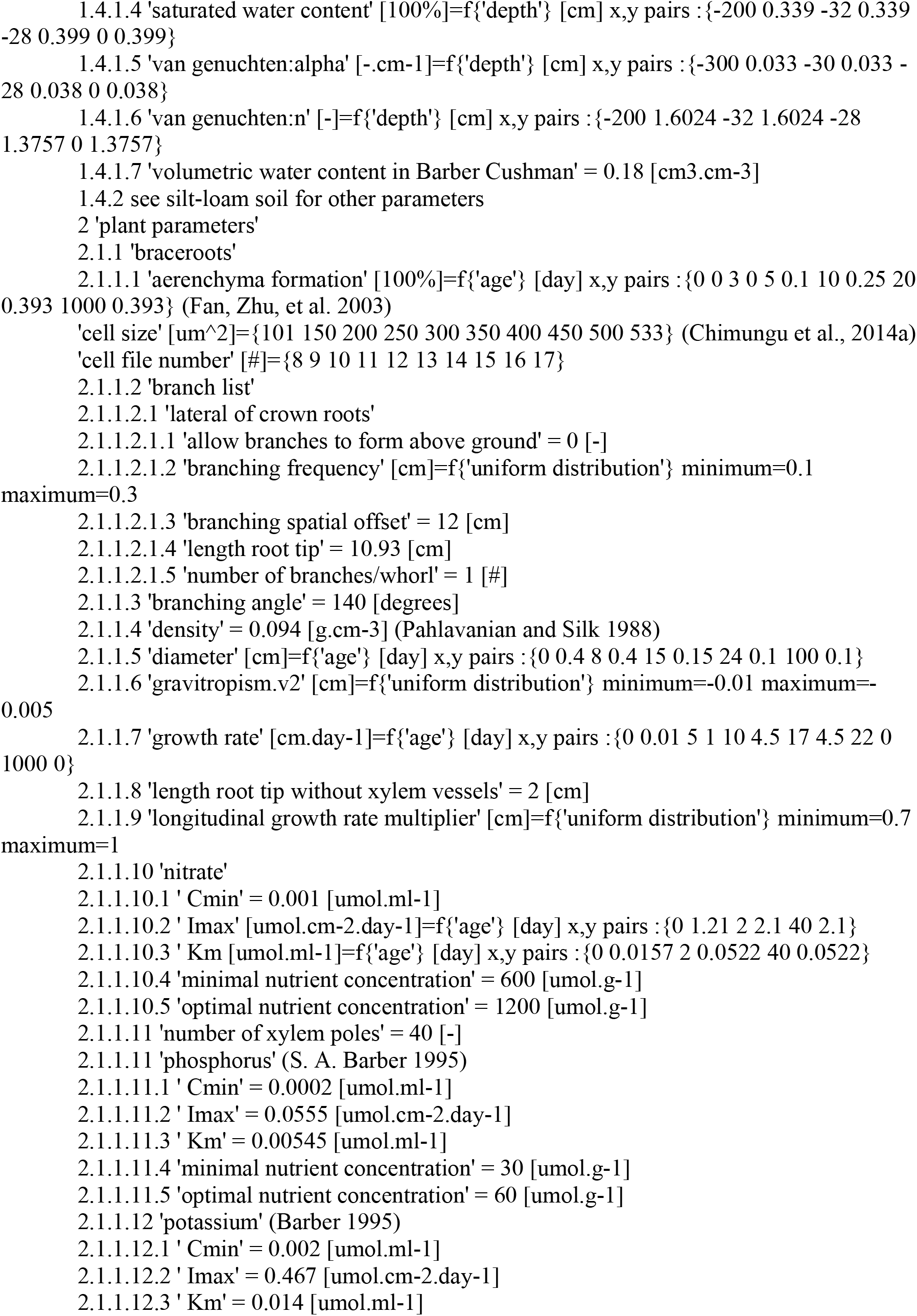
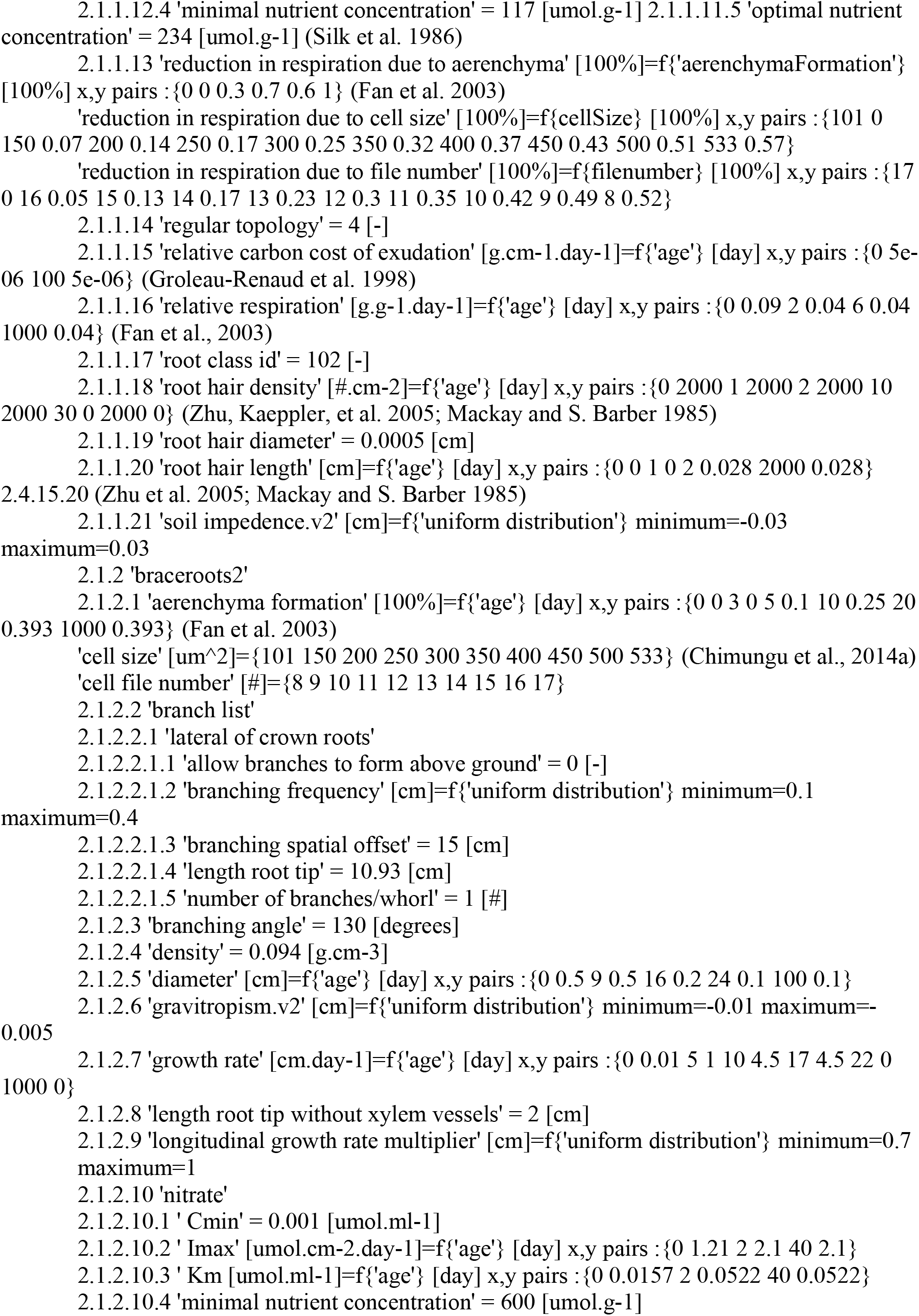
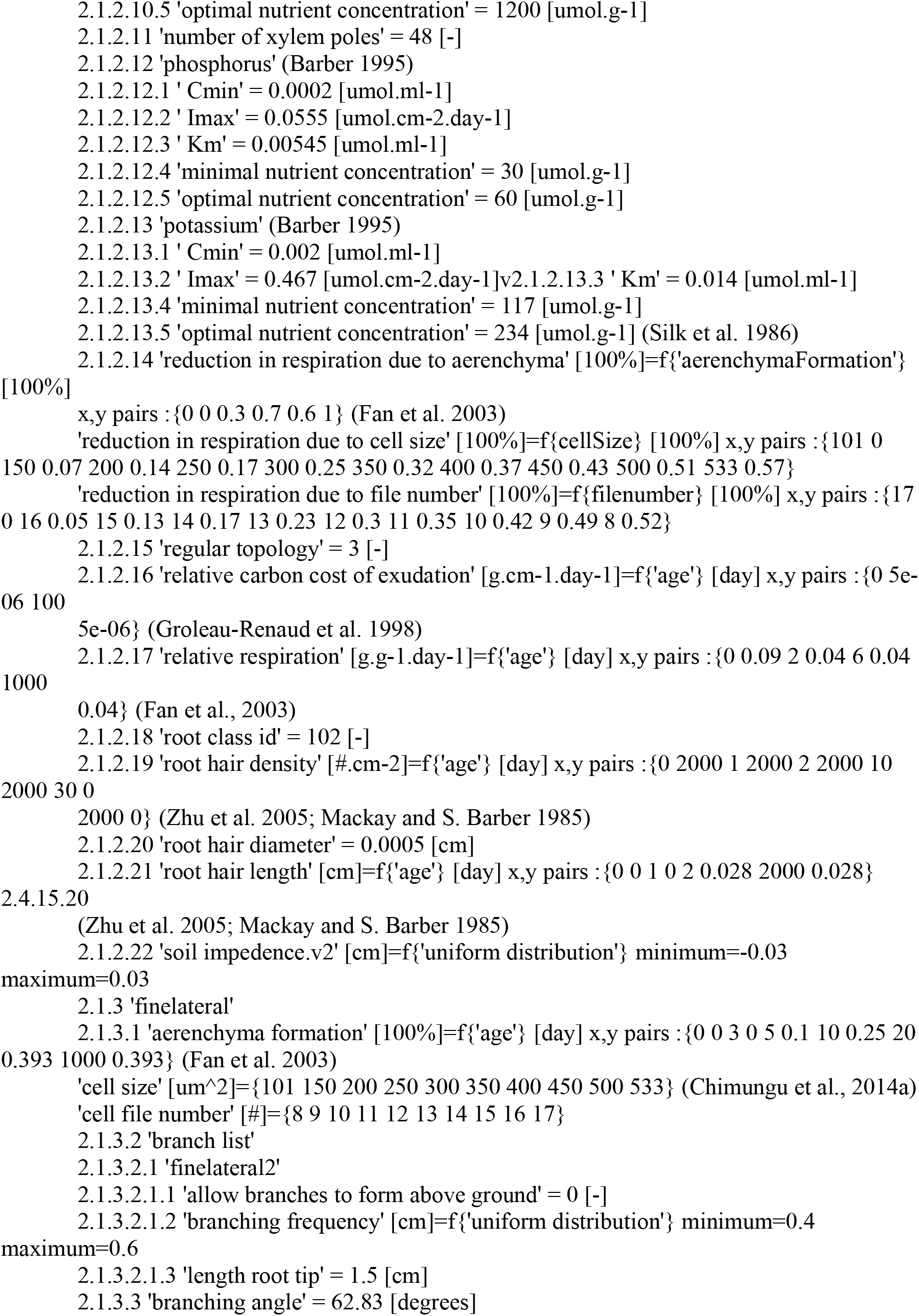
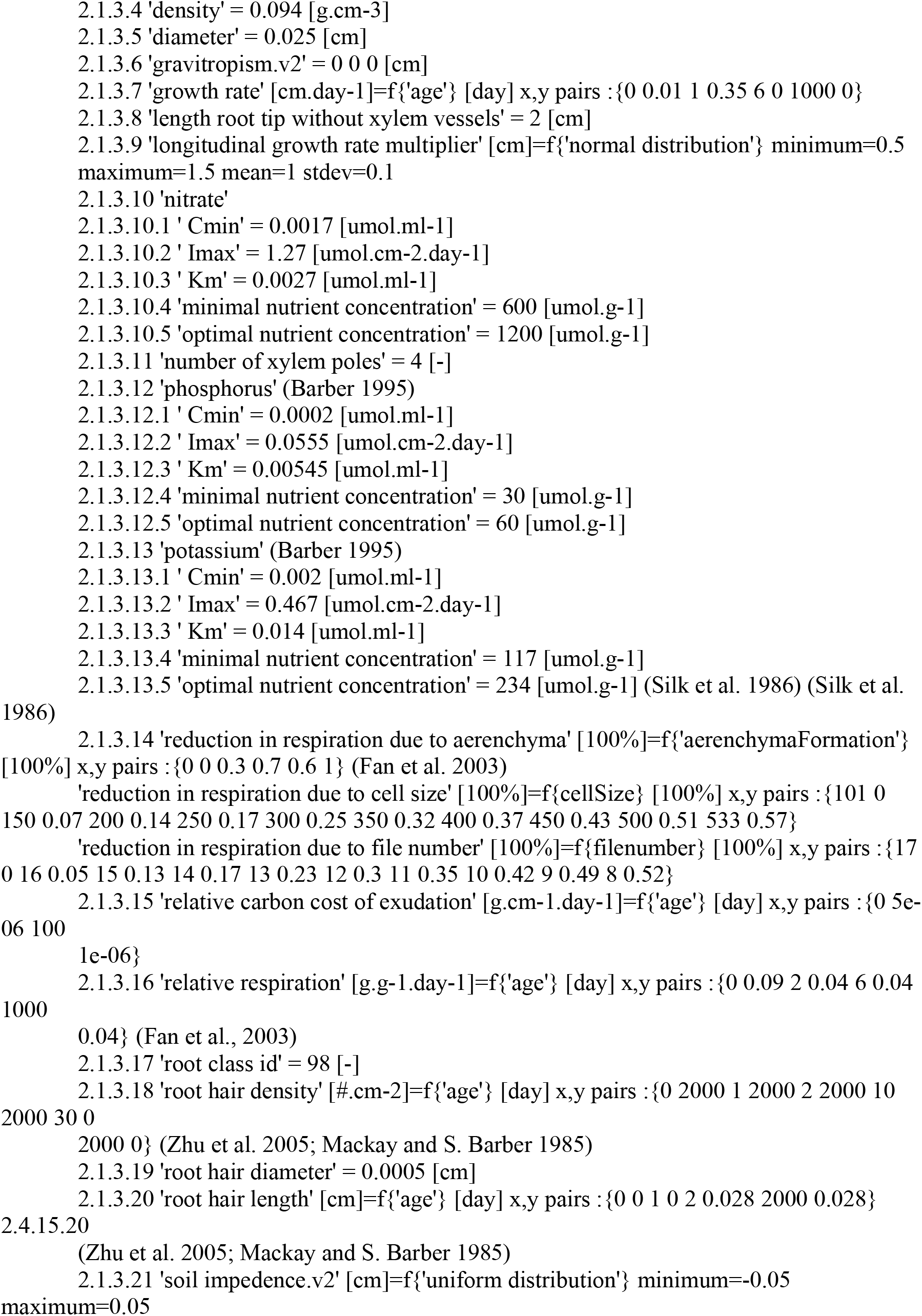
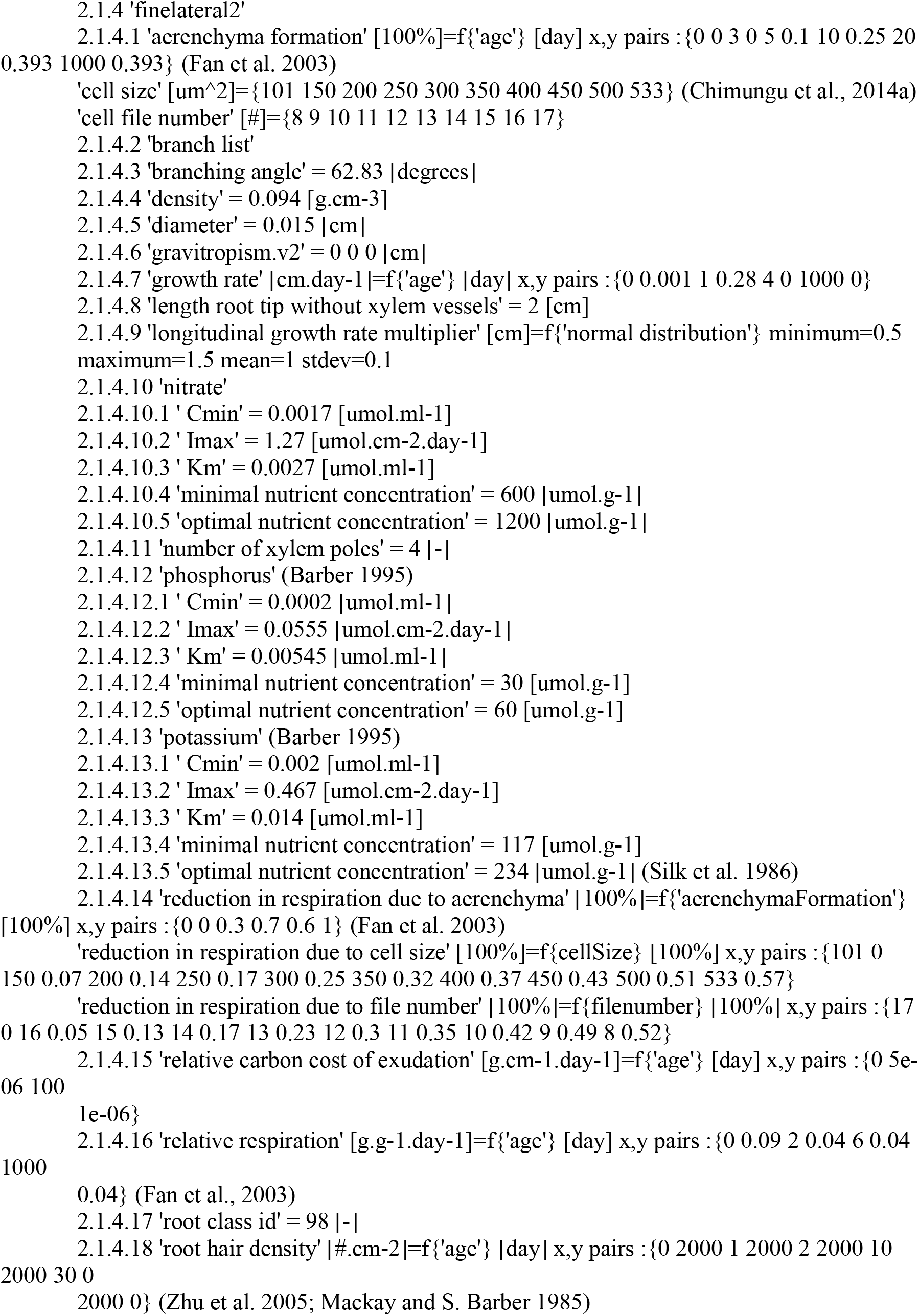
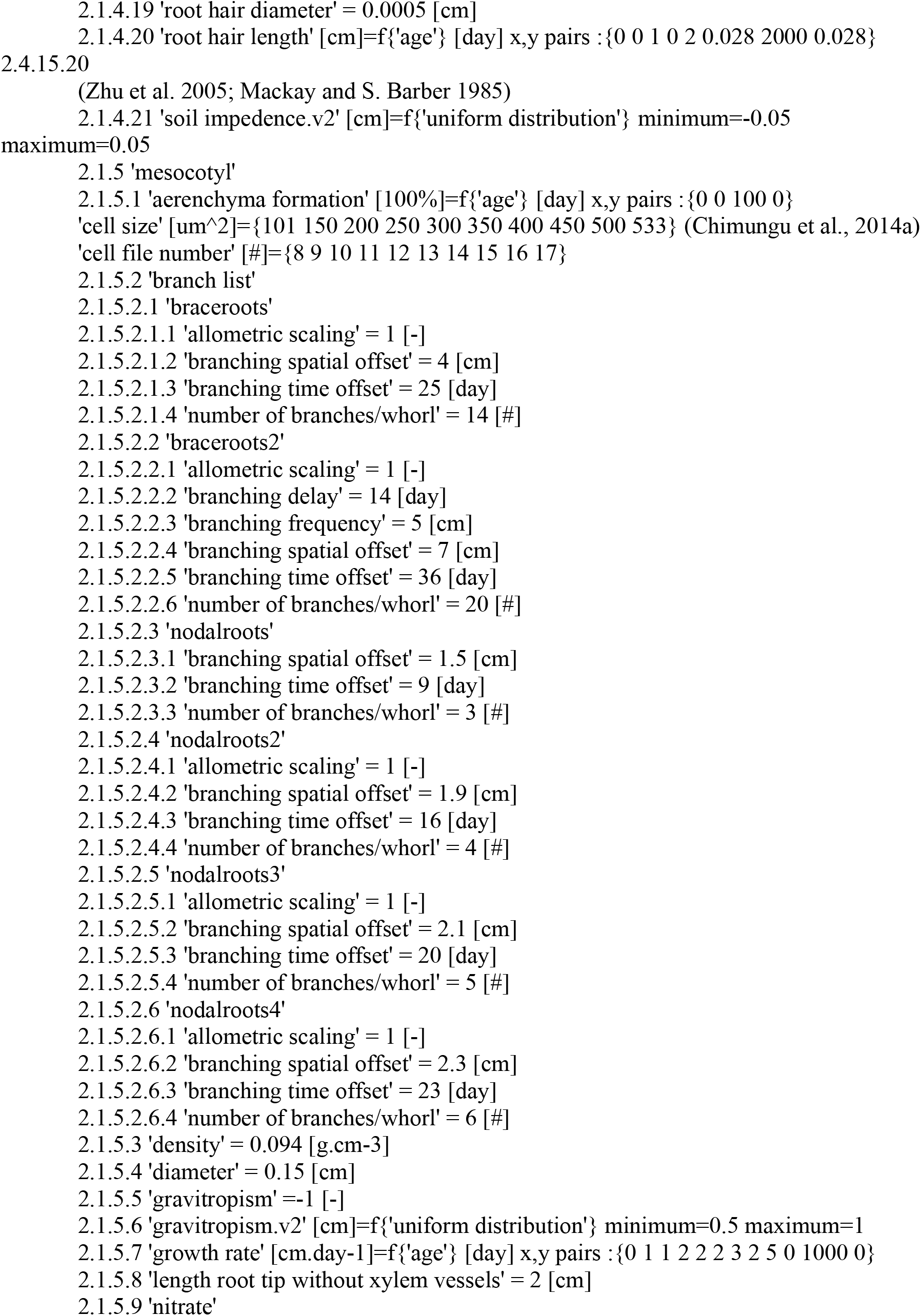
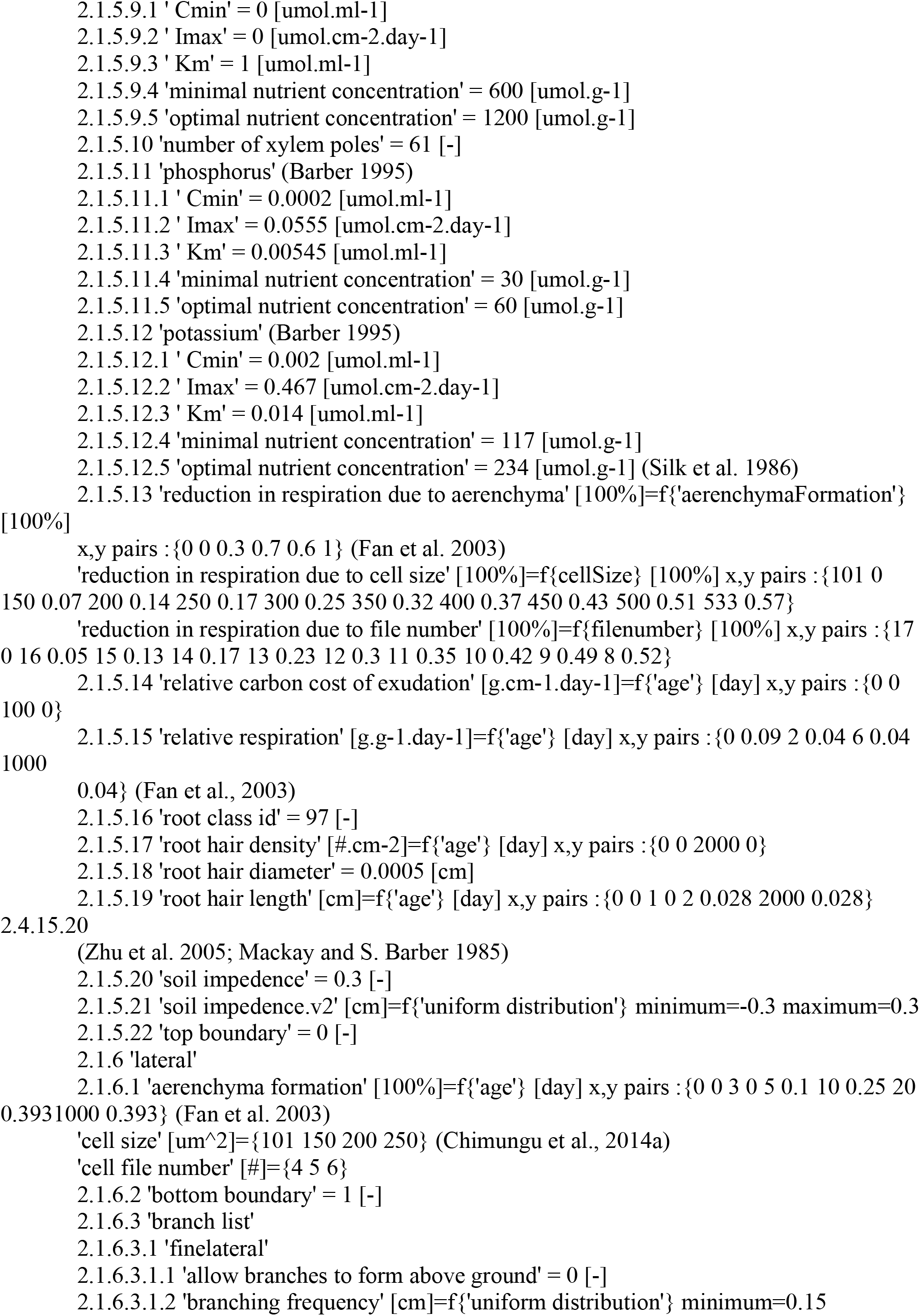
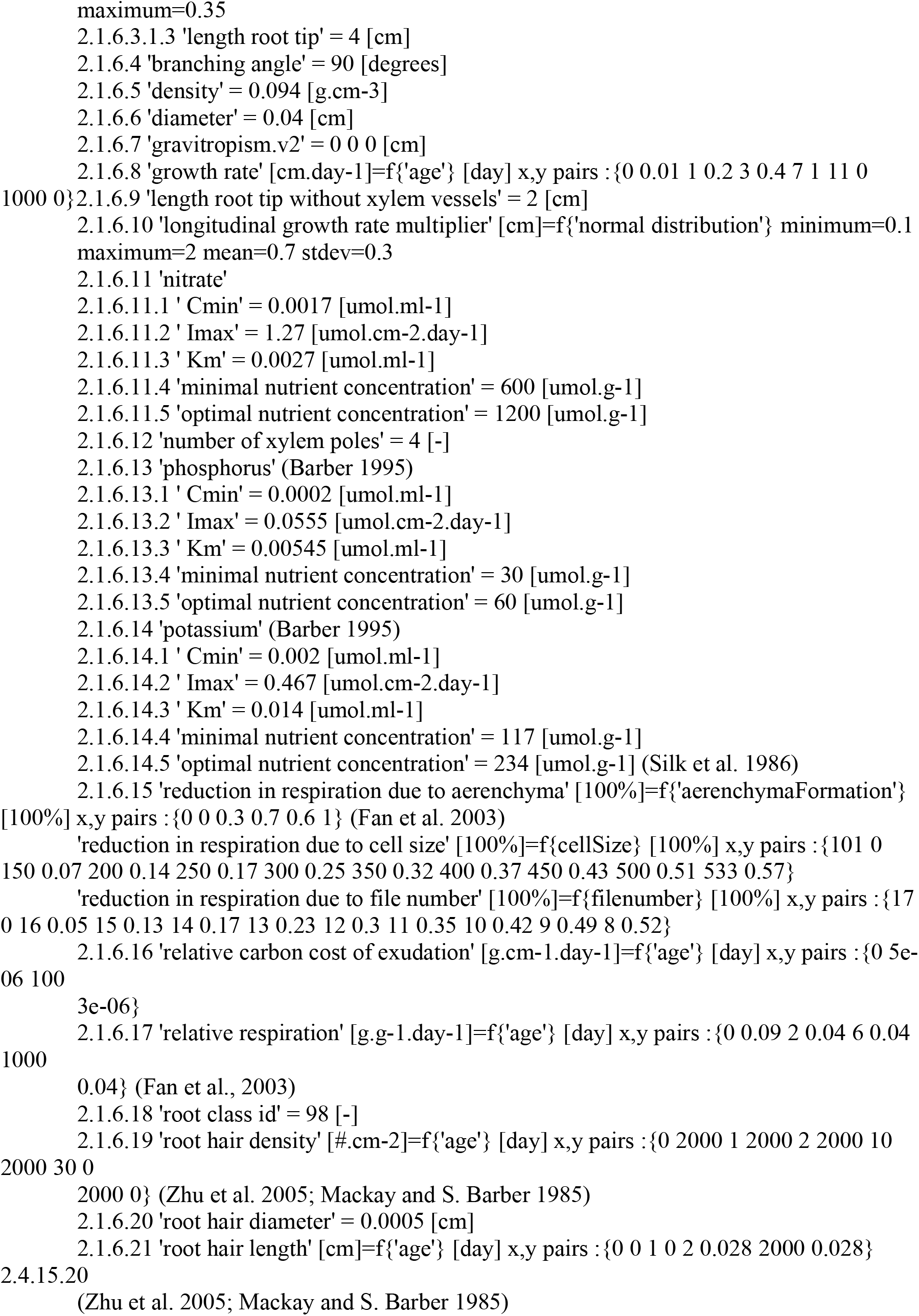
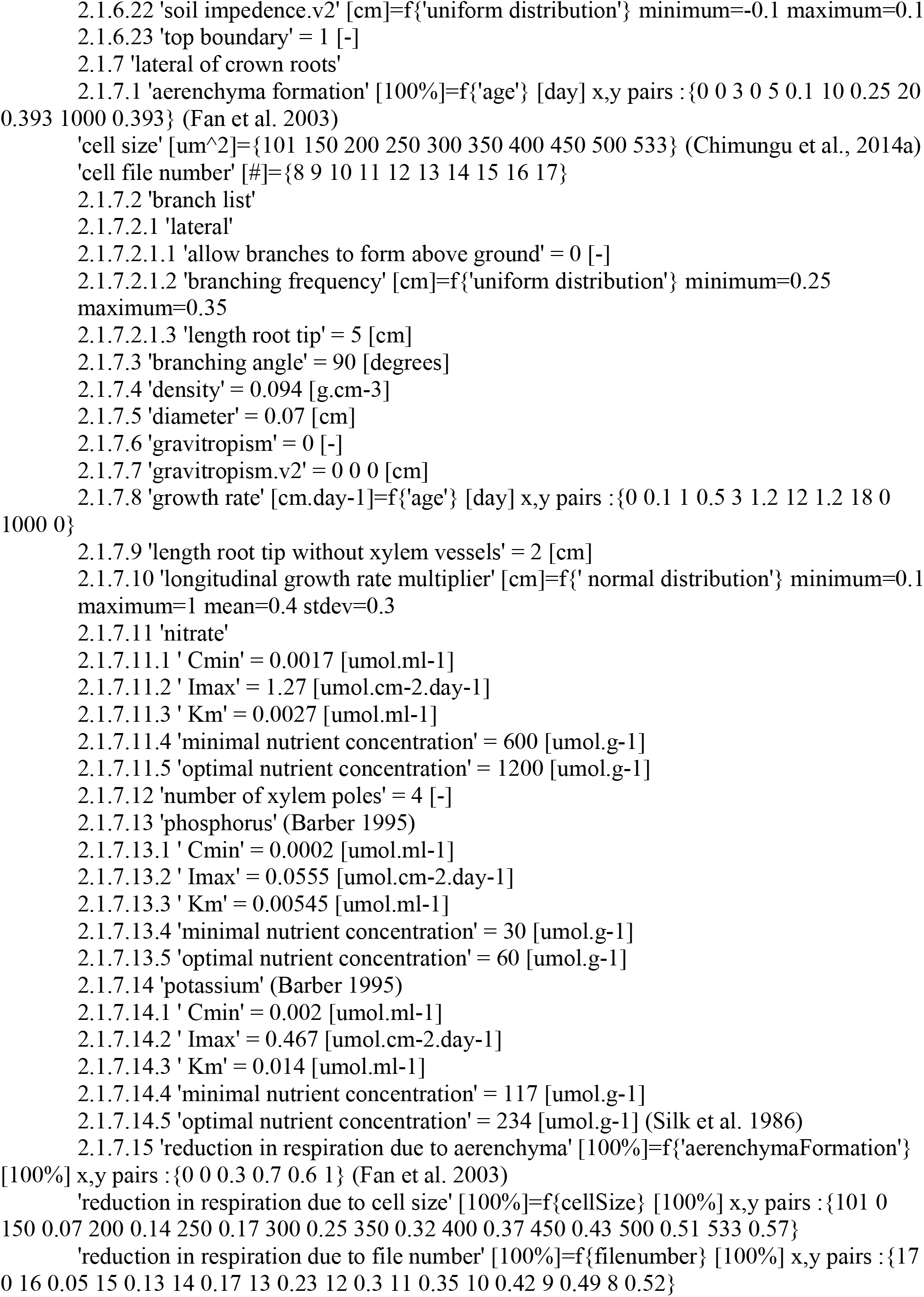
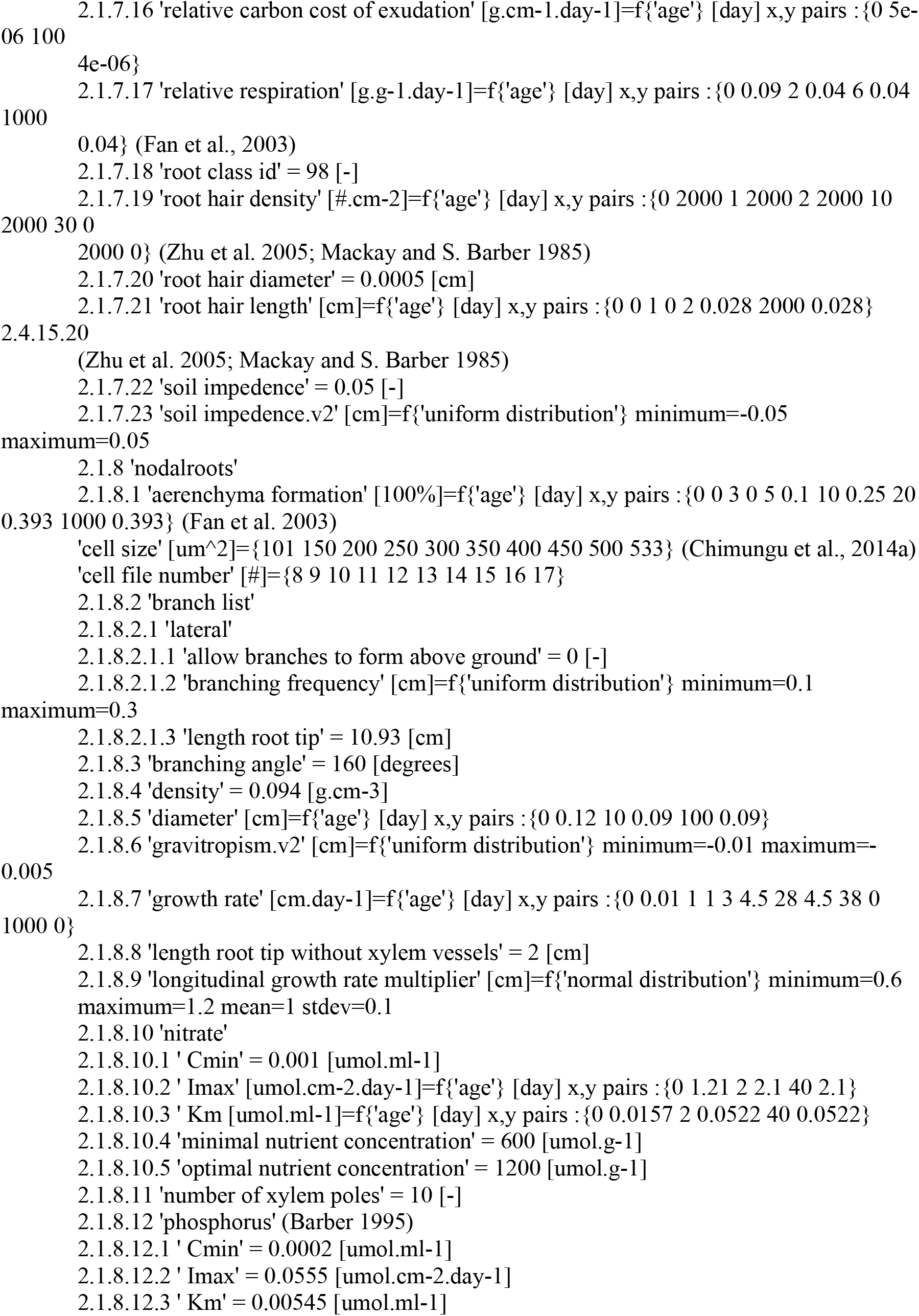
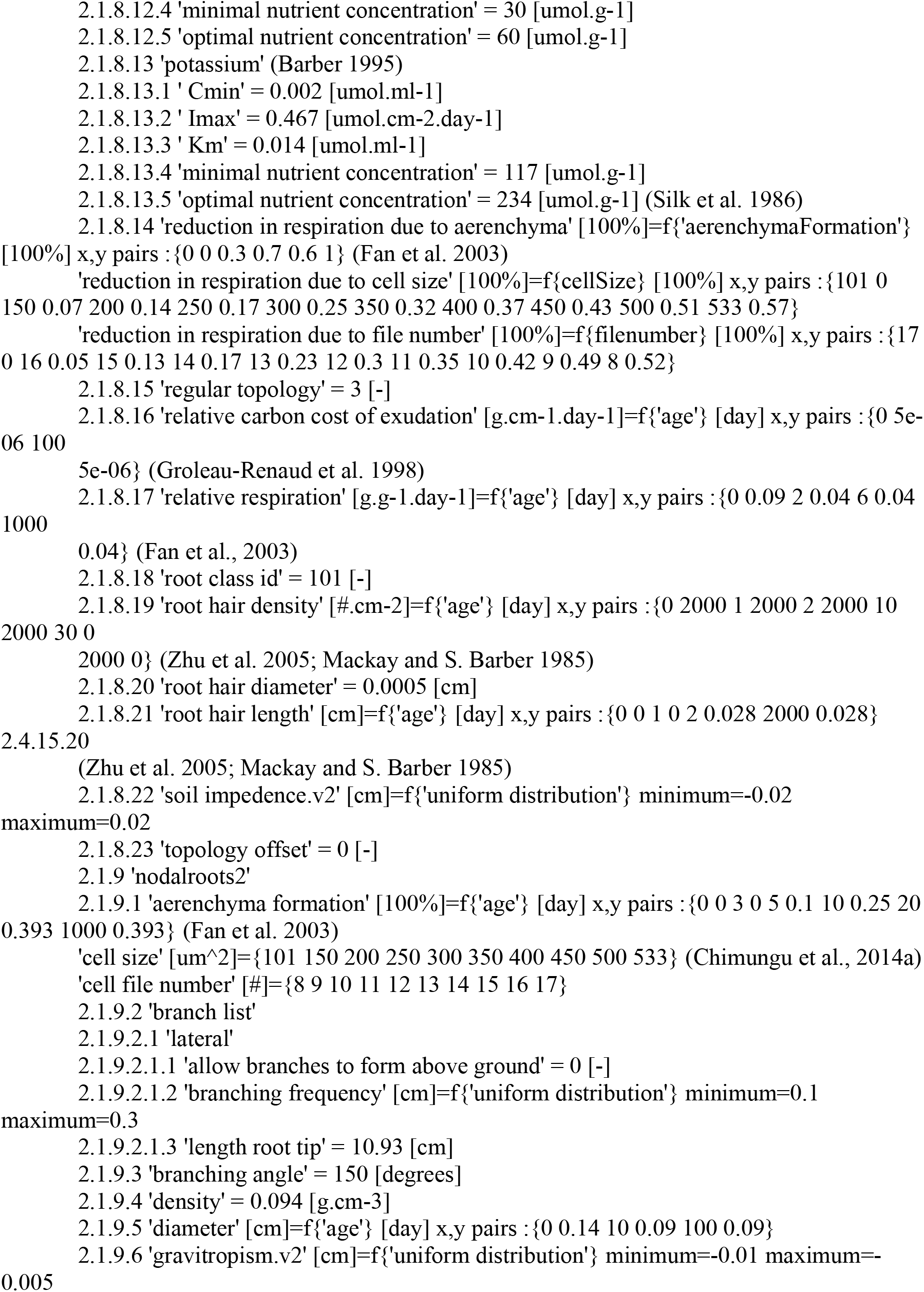
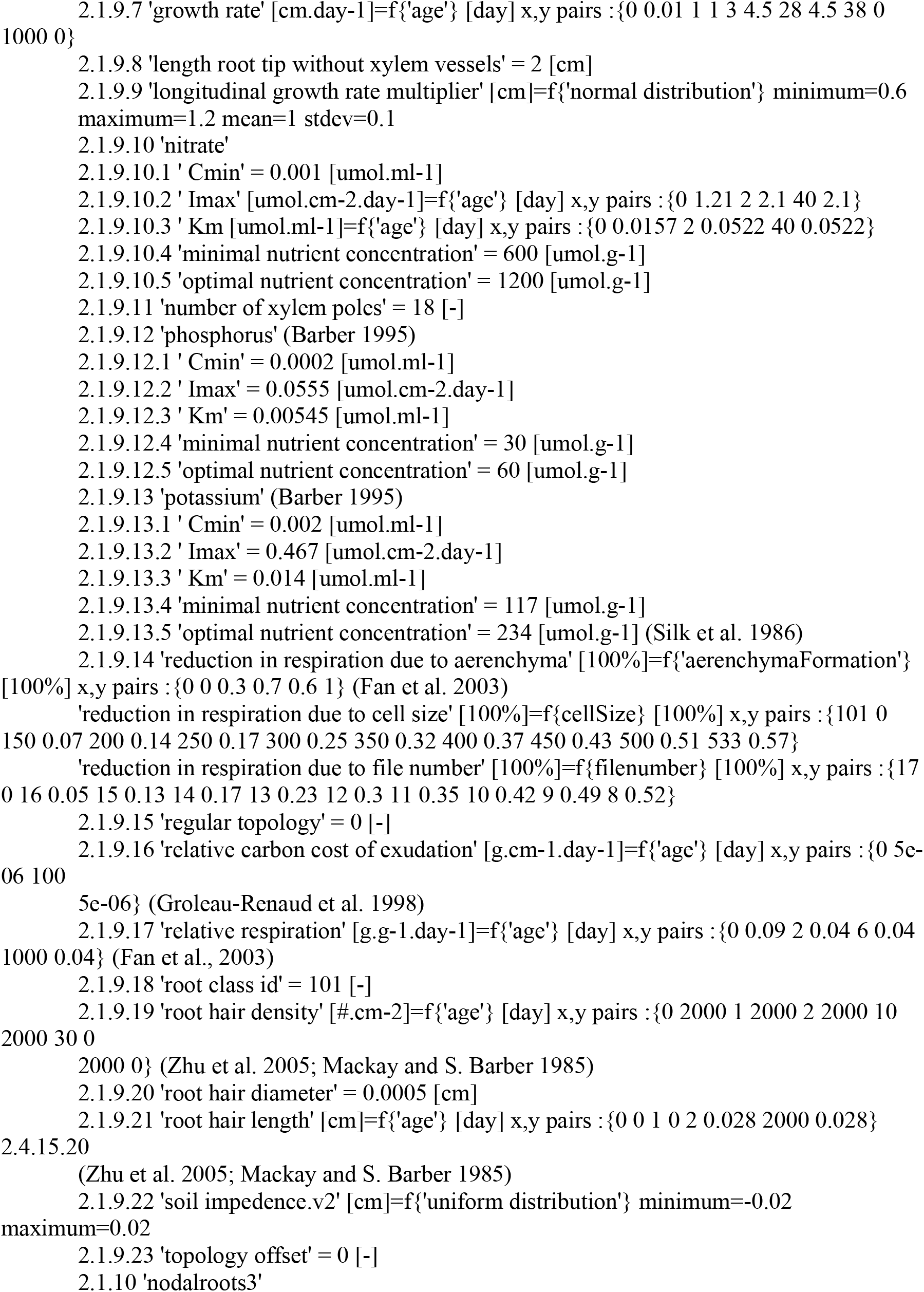
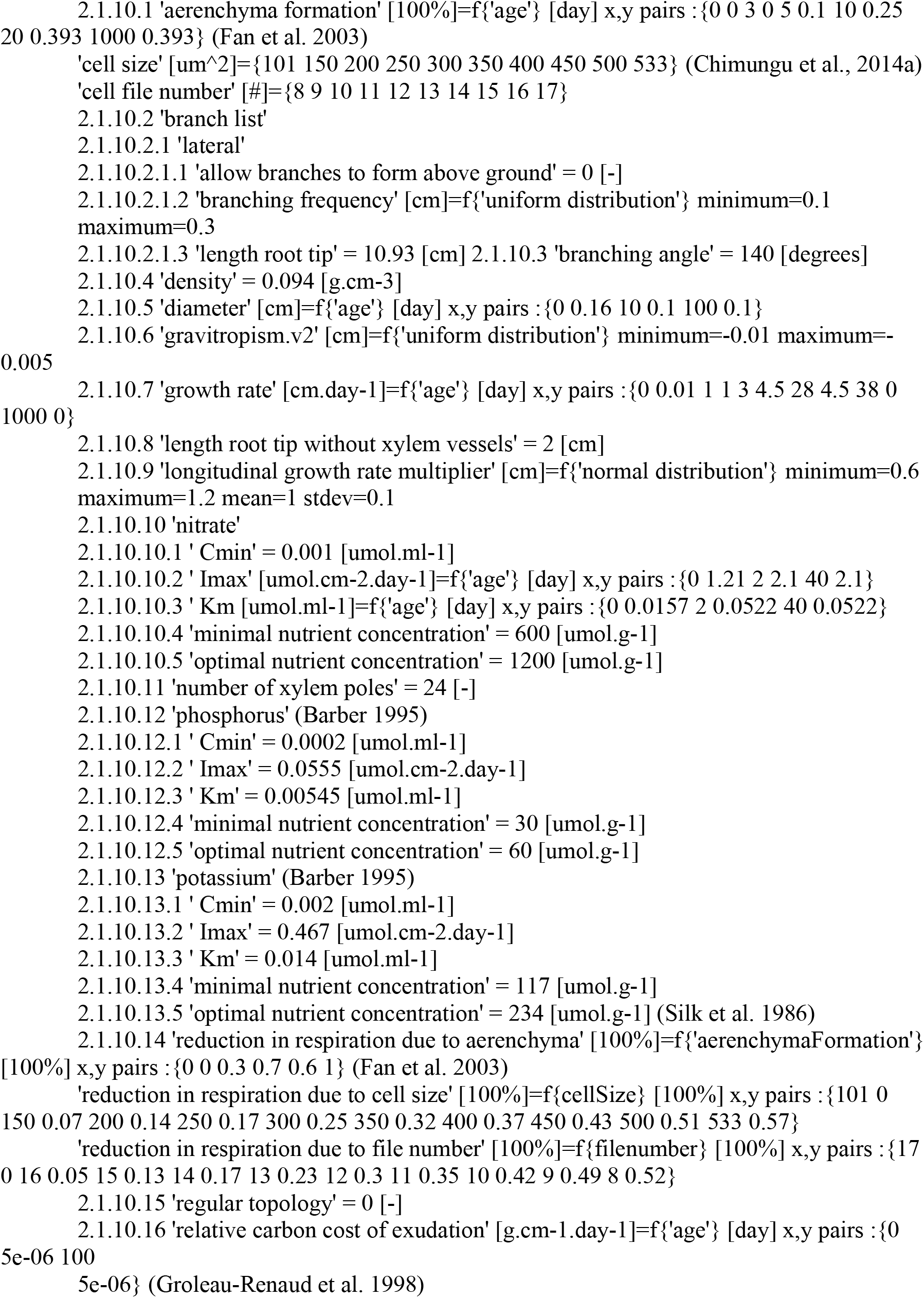
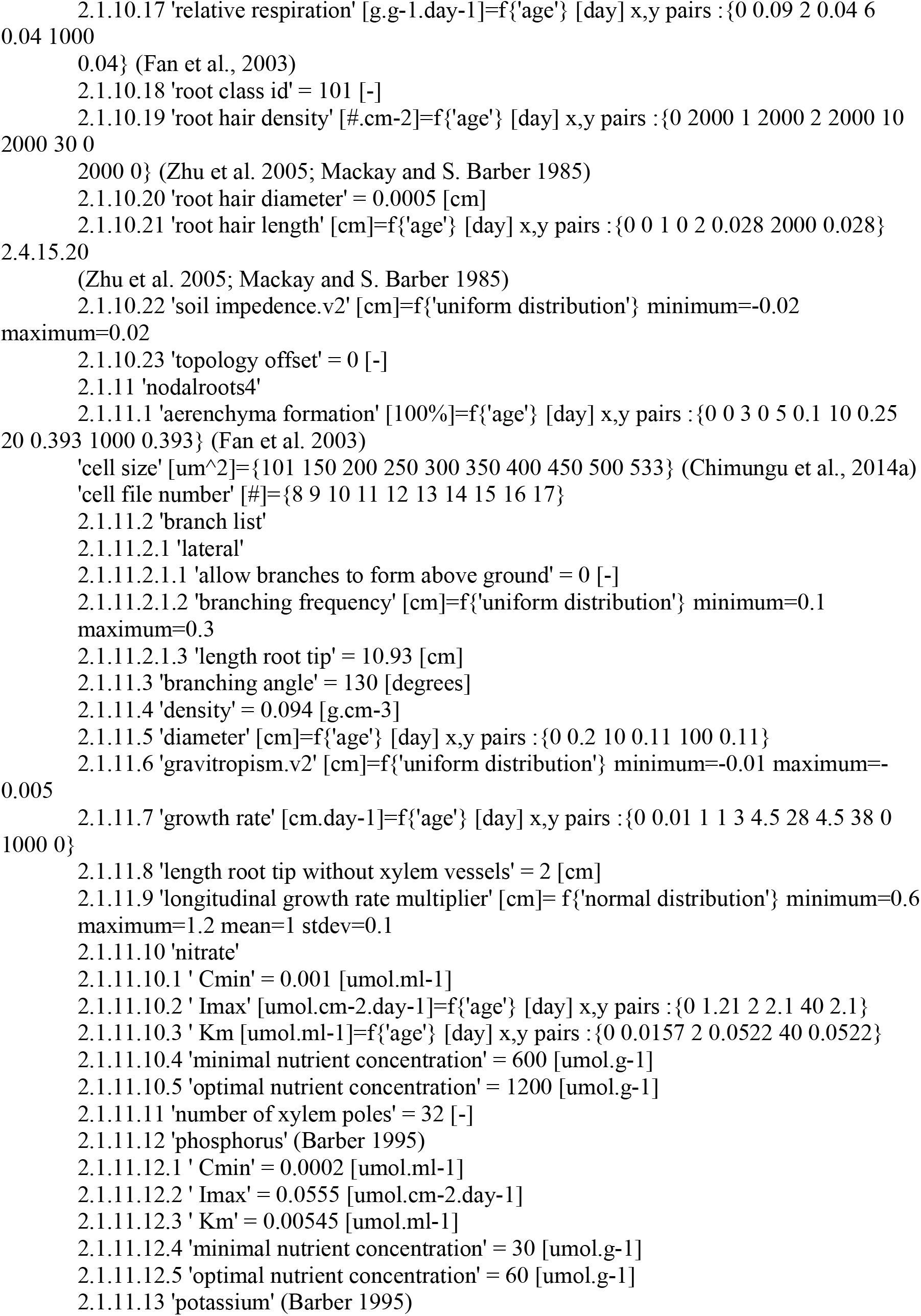
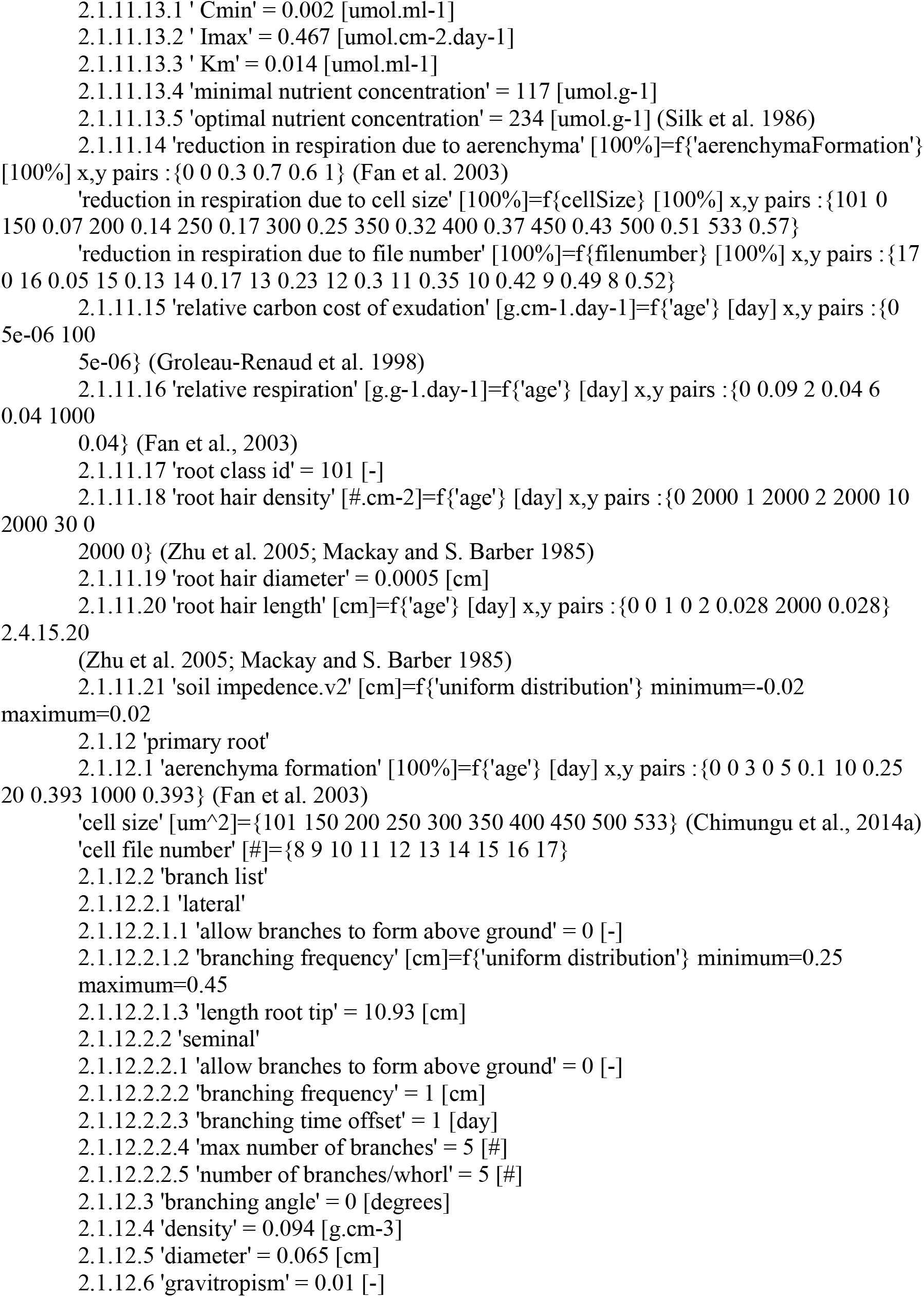
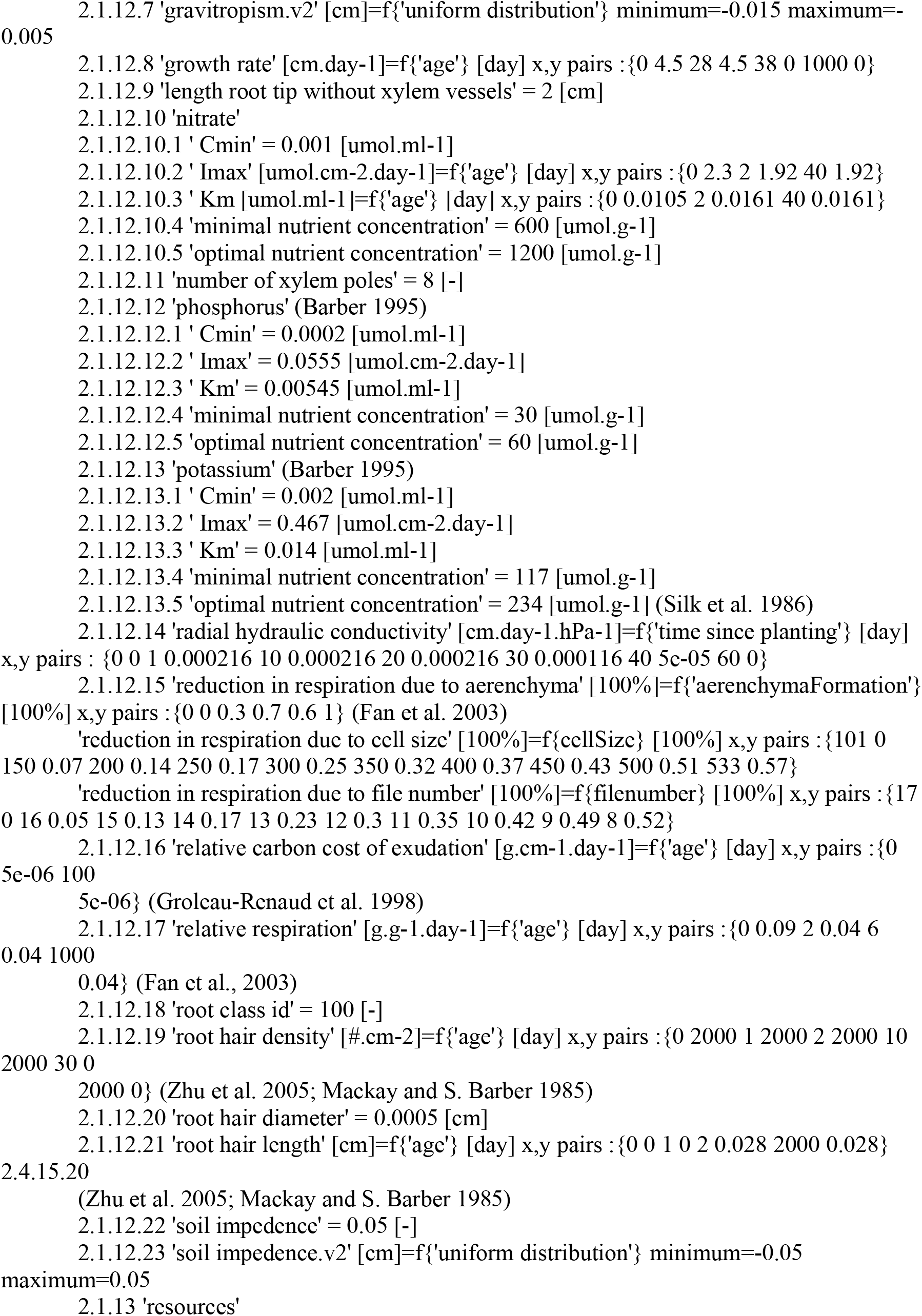
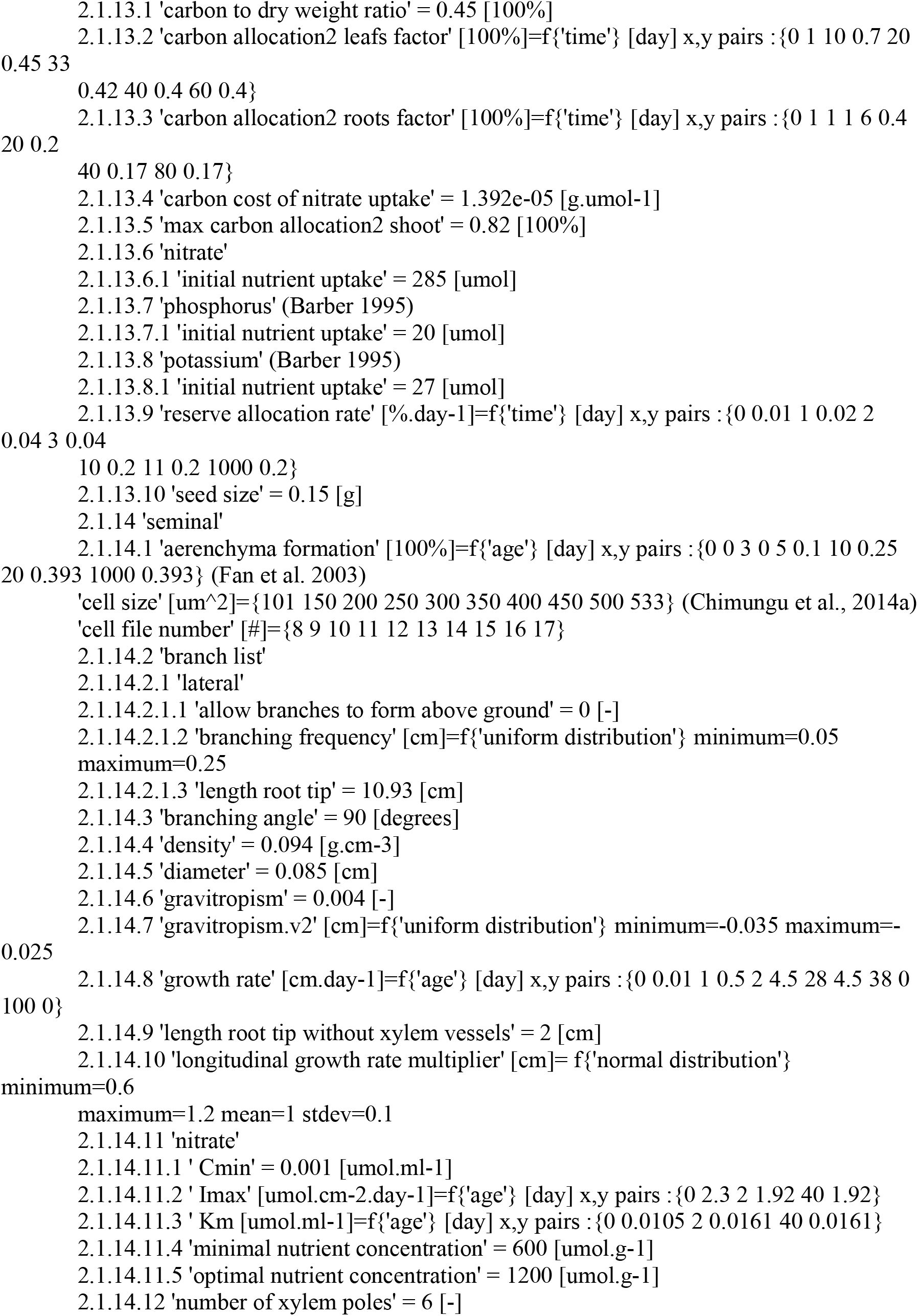
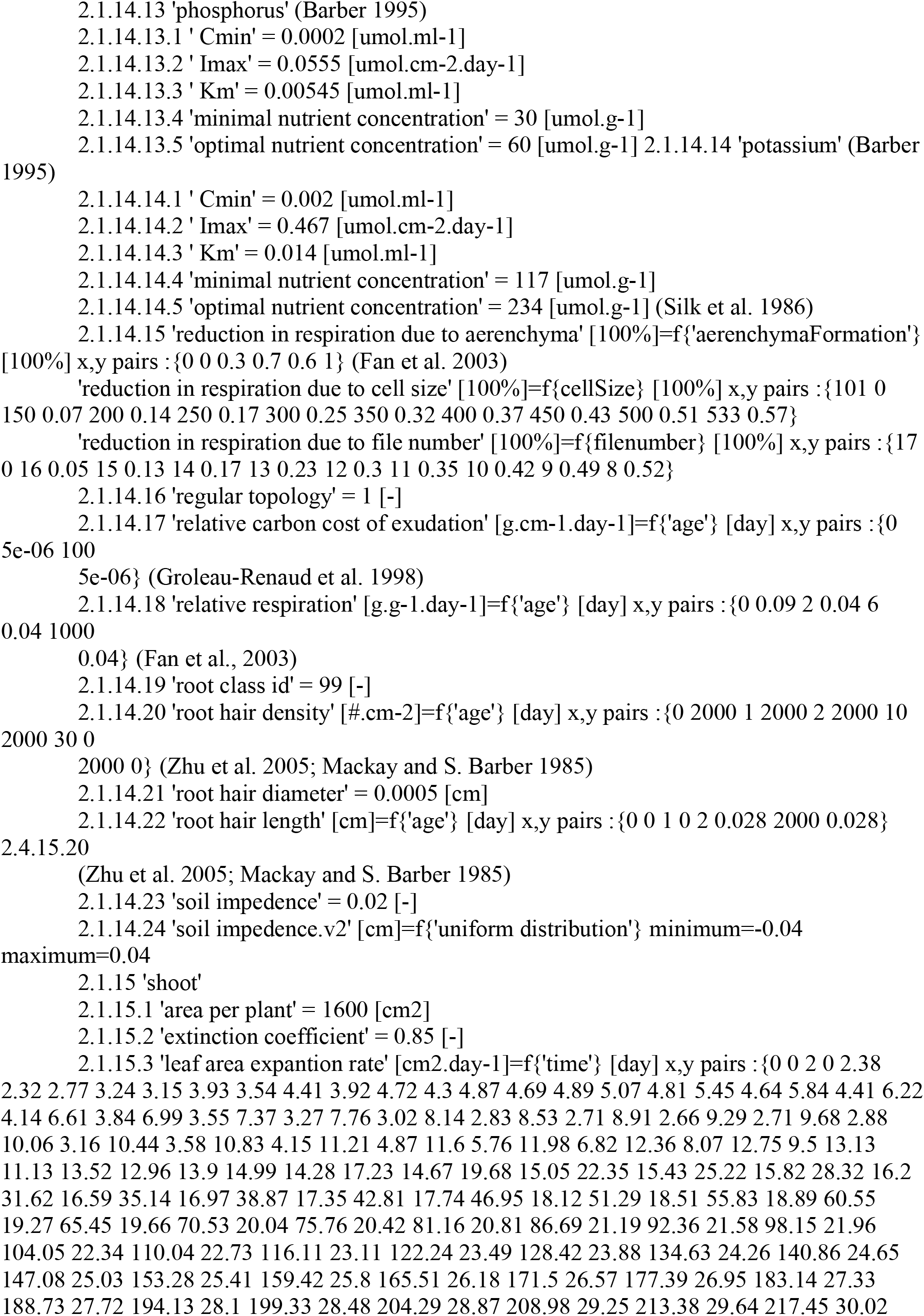
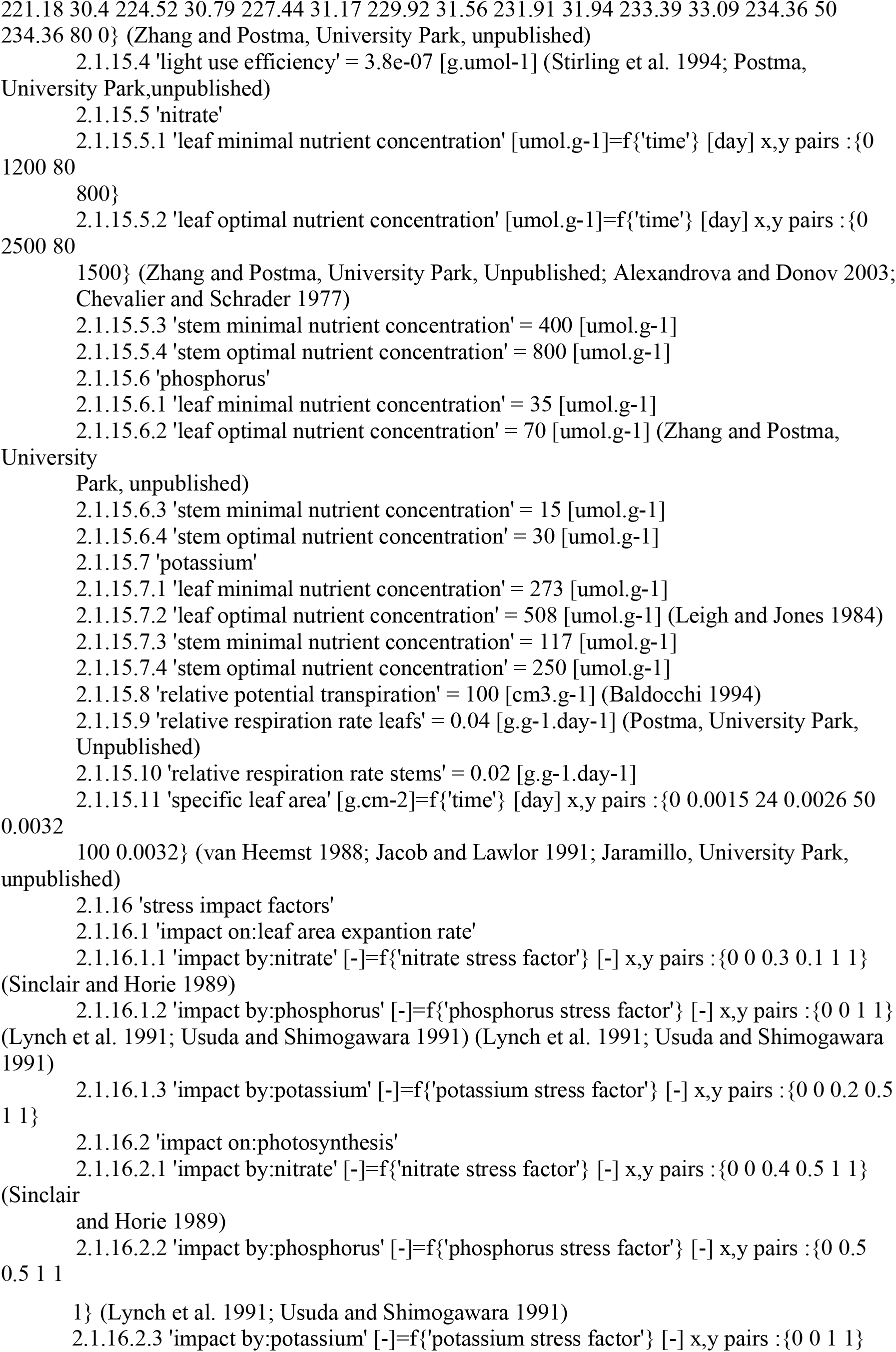

